# Overlap in synaptic neurological condition susceptibility pathways and the neural pannexin 1 interactome revealed by bioinformatics analyses

**DOI:** 10.1101/801563

**Authors:** Simona D Frederiksen, Leigh E Wicki-Stordeur, Leigh Anne Swayne

## Abstract

Many neurological conditions exhibit synaptic impairments, suggesting mechanistic convergence. Additionally, the pannexin 1 (PANX1) channel and signalling scaffold is linked to several of these neurological conditions and is an emerging regulator of synaptic development and plasticity; however, its synaptic pathogenic contributions are relatively unexplored. To this end, we explored connections between synaptic neurodevelopmental disorder and neurodegenerative disease susceptibility genes discovered by genome-wide association studies (GWASs), and the neural PANX1 interactome (483 PANX1-interacting proteins identified from mouse Neuro2a cells). To identify shared susceptibility genes, we compared synaptic suggestive GWAS candidate genes amongst autism spectrum disorders, schizophrenia, Parkinson’s disease, and Alzheimer’s disease. Next, to further probe PANX1 signalling pathways at the synapse, we used bioinformatics tools to identify PANX1 interactome signalling pathways and protein-protein interaction clusters. To shed light on synaptic disease mechanisms potentially linking PANX1 and these four neurological conditions, we performed additional cross-analyses between gene ontologies enriched for the PANX1 synaptic and disease-susceptibility gene sets. Finally, to explore the regional specificity of synaptic PANX1-neurological conditions connections, we identified brain region-specific elevations of synaptic PANX1 interactome and GWAS candidate gene set transcripts. Our results confirm considerable overlap in risk genes for autism spectrum disorders and schizophrenia and identify potential commonalities in genetic susceptibility for neurodevelopmental disorders and neurodegenerative diseases. Our findings also pinpointed novel putative PANX1 links to synaptic disease-associated pathways, such as regulation of vesicular trafficking and proteostasis, warranting further validation.

## INTRODUCTION

Dendritic spines are the site of post-synaptic communication between neurons. Several neurodevelopmental and neurodegenerative conditions, such as autism spectrum disorders (ASD), schizophrenia, Parkinson’s disease, and Alzheimer’s disease, exhibit divergent dendritic spine size, stability and/or function [1–8]. These alterations often precede obvious clinical symptoms, suggesting they could be involved in disease susceptibility and progression. Despite this, current understanding of the mechanisms affecting dendritic spine dynamics in these conditions is limited.

The pannexin 1 (PANX1) channel and cytoskeleton-regulating protein is emerging as a key regulator of dendritic spines. PANX1, highly expressed at post-synaptic densities [9], oligomerizes to form ion and metabolite channels, and also acts as a channel-independent signalling hub [10]. For example, we discovered protein-protein interactions (PPIs) between PANX1 and key cytoskeleton-regulating proteins involved in dendritic spine formation and stability [11–13], including collapsin response mediator protein 2 (CRMP2) and actin-related protein 3 (ARP3 of the ARP2/3 complex)[14–18]. As follows, PANX1 knock out mice exhibit several synaptic and behavioral abnormalities, such as altered hippocampal long-term potentiation and long-term depression, impaired object recognition, spatial memory and reversal learning, and increased anxiety [19–21]. Consistent with these findings, we discovered that PANX1 limits cortical neuron network size and complexity through inhibition of dendritic spine density and stability [22,23], and similarly, others have shown that PANX1 hinders hippocampal neuron spine maturation [24]. Not surprisingly, PANX1 and/or its protein interaction partners CRMP2 and ARP2/3 are implicated in several neurological conditions associated with dendritic spine abnormalities [25–31], raising the possibility that PANX1 and its interaction partners may contribute to disease risk and/or progression via dendritic spine modulation.

Therefore, here we aimed to identify potential links between synaptic neurological condition susceptibility genes and PANX1 by exploring the extent of overlap in (1) synaptic genes involved in four common neurological conditions associated with broad genetic susceptibility [32–36] and atypical dendritic spines, and (2) these same genes with the PANX1 interactome. The neurological conditions we selected, ASD, schizophrenia, Parkinson’s disease and Alzheimer’s disease, are commonly known and contribute substantial socioeconomic disease burden [37–40]. Moreover, we performed an in-depth bioinformatics analysis of the neural PANX1 interactome to identify PANX1 interaction partners potentially relevant to dendritic spine dynamics using *in silico* tools (Fig. 1). We first cross-referenced findings from Genome-Wide Association Studies (GWASs) for ASD, schizophrenia, Parkinson’s disease, and Alzheimer’s disease, to identify common suggestive causative genes. We then performed enrichment analyses of the total neural PANX1 interactome to identify overrepresented biological pathways (*e.g.,* relating to neurological diseases), including implicated PANX1-interacting proteins. We next identified links between existing PPI networks and synaptic PANX1-interacting proteins, the latter obtained by identifying protein hits from our mouse Neuro2a cell PANX1-EGFP interactome annotated to the Gene Ontology (GO) term ‘synapse’. This was done to gain further insight into the molecular mechanisms that might underly PANX1 regulation of dendritic spines. To investigate potential links between PANX1 and neurological conditions exhibiting dendritic spine pathology, we compared cellular localizations and biological functions of the synaptic PANX1 interactome with synaptic-enriched susceptibility genes for ASD, schizophrenia, Parkinson’s disease, and Alzheimer’s disease identified by GWASs. The outcomes of this work provide potential new insights into the role of PANX1 in the central nervous system (CNS) and suggest links between PANX1 and neurological conditions.

**Fig. 1:**
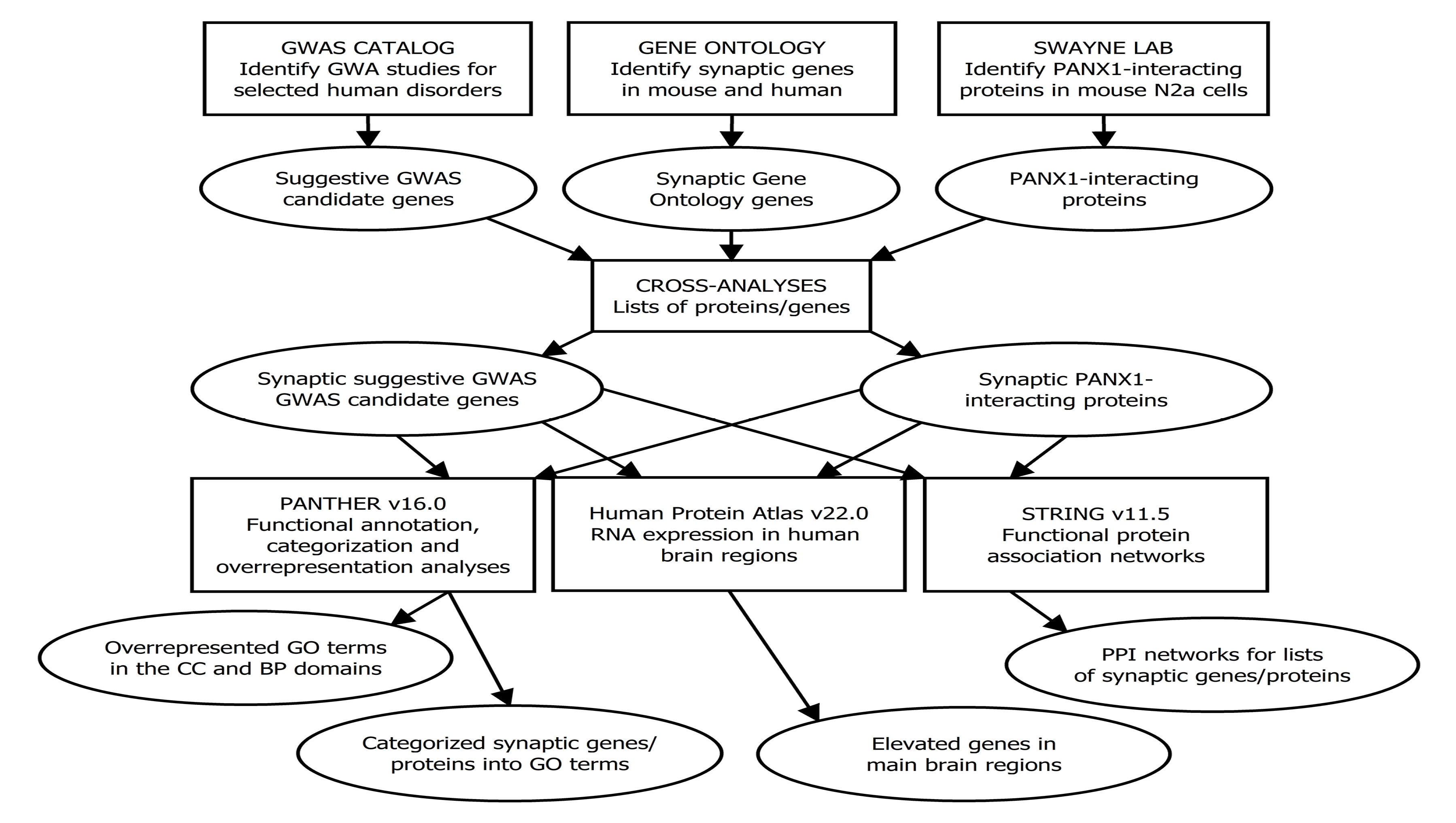
Work flow for the current study from materials and methods to results. In this study, a number of subanalyses were conducted, which explored the neurodevelopmental disorder and neurodegenerative disease susceptibility genes, and the PANX1 interactome in mouse N2a cells overlapping with the Gene Ontology (GO) synapse. The data was obtained from three sources: The genome-wide association study (GWAS) catalog [42], Swayne lab [our lab], and GO knowledge base via PANTHER [Protein ANalysis THrough Evolutionary Relationships] [49,56], which is a classification system. The data were analyzed using the statistical computing environment R and the following data or databases: PANTHER [Protein ANalysis THrough Evolutionary Relationships] [47–49], the Human Protein Atlas) [66,67] and STRING [Search Tool for the Retrieval of Interacting Genes/Proteins] [59–61]. In addition to these subanalyses, the findings across analyses were compared to provide a comprehensive overview for each neurodevelopment disorder and neurodegenerative disease in relation to the PANX1 interactome, and overrepresented PANTHER pathways were identified for the PANX1 interactome. *Abbreviations: BP, biological process; CC, cellular component*.

## MATERIALS AND METHODS

An overview of the study workflow can be found in Figure 1. For the comparisons and analysis, the R statistical computing environment v4.2.2 was applied. On June 2, 2020 the University of Victoria’s Human Research Ethics Board exempted the study from ethical review as the study: 1) is limited to accessing publicly available data sets; and 2) does not involve human participants. Biosafety approval was obtained from the University of Victoria Biosafety committee to undertake the experiments identifying PANX1 interacting proteins.

### DATA INPUTS

#### Extraction of suggestive GWAS candidate genes involved in neurodegenerative diseases and neurodevelopmental disorders in humans

On 13/11/2022, we searched for the Experimental Factor Ontology (EFO) [41] trait labels ‘autism spectrum disorder’, ‘schizophrenia’, ‘parkinson disease’, ‘alzheimer disease’ and separately in the NHGRI-EBI GWAS catalog API [42,43] by means of the gwasrapidd R package [44] and extracted information about the studies (some publications contain multiple GWASs), variants (or single nucleotide polymorphisms (SNPs)) and associations (SNP-trait associations). For Parkinson’s disease, 61 studies and 494 associations were available; for Alzheimer’s disease, 117 studies and 1988 associations were available; for schizophrenia, 131 studies, 4961 associations were available; and for ASD, 35 studies and 1275 associations were available. Information about the original publications, including accession ID of the GWAS Catalog study, can be found in Table S1.

Using the extracted data, associations without gene information were excluded (gene symbols or entrez IDs; symbols beginning with LOC were kept) and only suggestive GWAS candidate genes reported more than once were included for further analysis. Based on these filtering criteria, a total of 461, 881, 74 and 86 suggestive GWAS candidate genes were identified for ASD, schizophrenia, Parkinson’s disease, and Alzheimer’s disease, respectively. The complete lists of suggestive GWAS candidate genes obtained after filtering were used as inputs for the cross-analyses.

#### Identification of the PANX1 interactome in mouse N2a cells

We previously identified the putative PANX1 interactome from mouse N2a neuroblastoma-derived cells, using methods that were comprehensively described in that work [11,13]. Briefly, proteins co-precipitating with PANX1EGFP [enhanced green fluorescent protein] or EGFP from N2a cells, were identified by the UVIC-Genome BC Proteomics Centre using high performance liquid chromatography-tandem mass spectrometry (LC-MS/MS) followed by analysis with Proteome Discoverer v1.3.0.339 (Thermo Scientific) and Mascot v2.2 [45] [percolator settings: Max delta Cn 0.05, Target false discovery rate (FDR) strict 0.01, Target FDR relaxed 0.05 with validation based on q-value]. The q-value refers to the minimal FDR at which a peptide spectrum match was accepted [46]. To identify proteins selectively interacting with PANX1, all proteins co-precipitating with anti-GFP antibody from EGFP-expressing cells were removed from the list of PANX1 interactors. This paradigm was repeated 3 times and the results were pooled. The complete list of PANX1-interacting proteins identified from those experiments (not previously published in its entirety) was used as input for the cross-analyses.

### CROSS-ANALYSIS

#### Cross-analysis of the suggestive disease susceptibility GWAS candidate genes in humans and PANX1 interactome in mice with the synapse GO term

To identify known synaptic genes, genes annotated to the synapse GO term were extracted for *Homo Sapiens* and *Mus Musculus* using the PANTHER database v17.0 and the GO Ontology database released on 01/07/2022 [47–51]. Second, the entrez IDs were obtained by converting UniProtKB IDs from the *Homo Sapiens* synapse GO term gene list using the UniProt Retrieve/ID mapping tool (https://www.uniprot.org/id-mapping), and finding the human orthologs for the MGI IDs from the *Mus Musculus* GO term gene list using the biological DataBase network (bioDBnet; http://biodbnet.abcc.ncifcrf.gov) [52]. When unable to obtain the entrez IDs, we manually looked up the entrez IDs in the HUGO Gene Nomenclature Committee (HGNC) database [53], or the human orthologs in the Mouse Genome Informatics (MGI) international database [54,55]. The gene lists were combined and used as input for the cross-analyses.

To allow for comparison with the GWAS findings, human orthologs in the form of entrez IDs were identified for the mouse UniProtKB IDs forming the PANX1 interactome using bioDBnet [52]. As described above, for the genes we were unable to obtain entrez IDs, we manually looked up the human orthologs in the MGI international database [54].

Next, we conducted cross-analyses between the GO term synapse (GO:0045202; identified using the PANTHER database [48,49,56]) and the (i) suggestive GWAS candidate genes involved in the selected human neurodegenerative diseases and neurodevelopmental disorders, and (ii) proteins comprising the mouse PANX1 interactome. This resulted in identification of synaptic genes/proteins, and subsequently overlaps between the lists of synaptic suggestive GWAS candidate genes for the neurological diseases and synaptic PANX1-interacting proteins were examined.

### DOWNSTREAM ANALYSES

#### Functional annotation, categorization and overrepresentation analyses of the synaptic suggestive GWAS candidate genes and PANX1 interactome using the PANTHER database

Bioinformatics analysis of the synaptic suggestive GWAS candidate genes for each neurodevelopmental disorder and neurodegenerative disease (using *Homo Sapiens* Entrez gene identifiers), proteins from our PANX1-interacting protein list (using *Mus Musculus* UniProtKB unique identifiers), and synaptic proteins from our PANX1-interacting protein list (using *Mus Musculus* UniProtKB unique identifiers or *Homo Sapiens* Entrez gene identifiers) was carried out using the curated database PANTHER v17.0 [48,49,56]. The genes/proteins were annotated to (i) PANTHER pathways [57] created using the CellDesigner tool, a modelling tool for biochemical networks [58], and/or (ii) GO terms within the Biological Process and Cellular Component domains from the GO knowledge base [50,51]. Right-tailed Fisher’s exact tests were used to identify overrepresented PANTHER pathways and GO terms (present in greater abundance than would be expected). FDR-corrected p-values < 0.05 (to account for multiple testing) were considered statistically significant. In addition, only PANTHER pathways and GO terms with at least 10 annotated proteins/genes were presented (to reduce the likelihood of false positives; to allow for comparison, this was not done for Table 5 and corresponding analysis).

### PPI network for the PANX1 synaptic interactome

The synaptic PPI network, based on interaction evidence [from STRING-defined categories: known interactions (curated databases, experimentally determined)], was created for *Mus Musculus* using the Search Tool for the Retrieval of Interacting Genes/Proteins (STRING) v11.5 [59–61], and the identified PANX1-interacting synaptic proteins were used as input (those proteins overlapping with the *Mus Musculus* synapse GO term using the PANTHER v17.0 database and the GO Ontology database released on 01/07/2022 [47–51]). Edges (also termed PPIs) were formed if the interaction score was at least 0.4 (medium confidence, *“estimated likelihood that a given interaction is biologically meaningful, specific and reproducible, given the supporting evidence”* [62]), and the thickness of the edges in the PPI network indicates the strength of data support and dashed line edges reveal PPIs between clusters. First, clusters were identified using the unsupervised Markov Cluster (MCL) algorithm with inflation factor 1.3 (higher inflation factor leads to more clusters, but noting that *“MCL is remarkably robust to graph alterations”* [63–65]. The outcome of this analysis was used to select the final number of clusters in the PPI network, and kmeans clustering was carried out. The potential function of the clusters (formed by at least 4 proteins) were investigated using the PANTHER database v17.0 [47–49] focusing specifically on PANTHER protein classes and GO biological process terms.

#### Synaptic PANX1-interacting proteins and suggestive GWAS candidate genes expressed in specific brain regions in humans

RNA expression in human brain regions was explored using the Human Protein Atlas v22.0 (<u>proteinatlas.org</u>) [66,67], and focusing specifically on genes classified as ‘regionally elevated’. The findings were compared with the synaptic PANX1 interactome and suggestive GWAS candidate genes for ASD, schizophrenia, Parkinson’s disease, and Alzheimer’s disease. Genes elevated in the following nine brain regions were included in the cross-analysis (UniProtKB IDs were converted to entrez IDs as described above): Cerebral cortex, hippocampal formation, amygdala, thalamus, hypothalamus, midbrain, pons, cerebellum, and medulla oblongata (https://www.proteinatlas.org/humanproteome/brain).

## RESULTS

### Comparison amongst neurological diseases exhibiting impaired synapse structure and/or stability revealed shared synaptic suggestive GWAS candidate genes

We focused our study on four major neurological conditions exhibiting synapse instability, namely ASD, schizophrenia, Parkinson’s disease, and Alzheimer’s disease. ASD and schizophrenia had more synaptic suggestive GWAS candidate genes in common than any other disease-disorder combination we studied (Figures 2-3; Table S2), consistent with other recent findings [68]. In fact, all the synaptic suggestive GWAS candidate genes identified for ASD overlapped with those identified for schizophrenia. These included well-known synaptic genes, such as those encoding for scaffold-protein ANKG (*ANK3*), the ionotropic glutamate receptor GluN2a (*GRIN2A*), and Ras GTPase activating protein 1 (*SYNGAP1*).

**Fig. 2.**
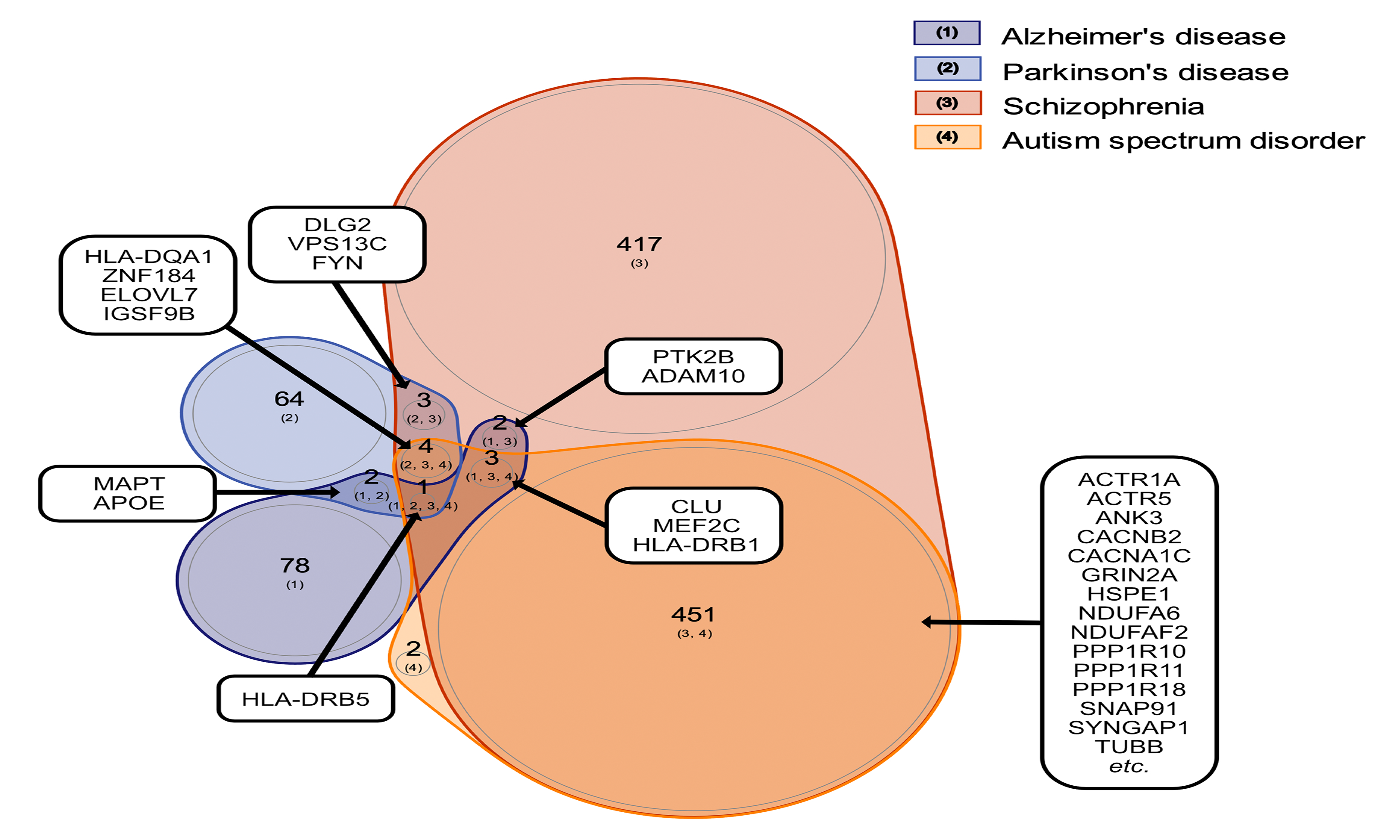
Overlap between genes associated with neurodevelopmental disorders and neurodegenerative diseases in humans based on genome-wide association study (GWAS) findings. Venn diagram displaying the number of suggestive candidate genes the selected neurodevelopmental disorders, autism spectrum disorder (ASD) and schizophrenia, and the selected neurodegenerative diseases, Parkinson’s disease and Alzheimer’s disease, have in common.

**Fig. 3.**
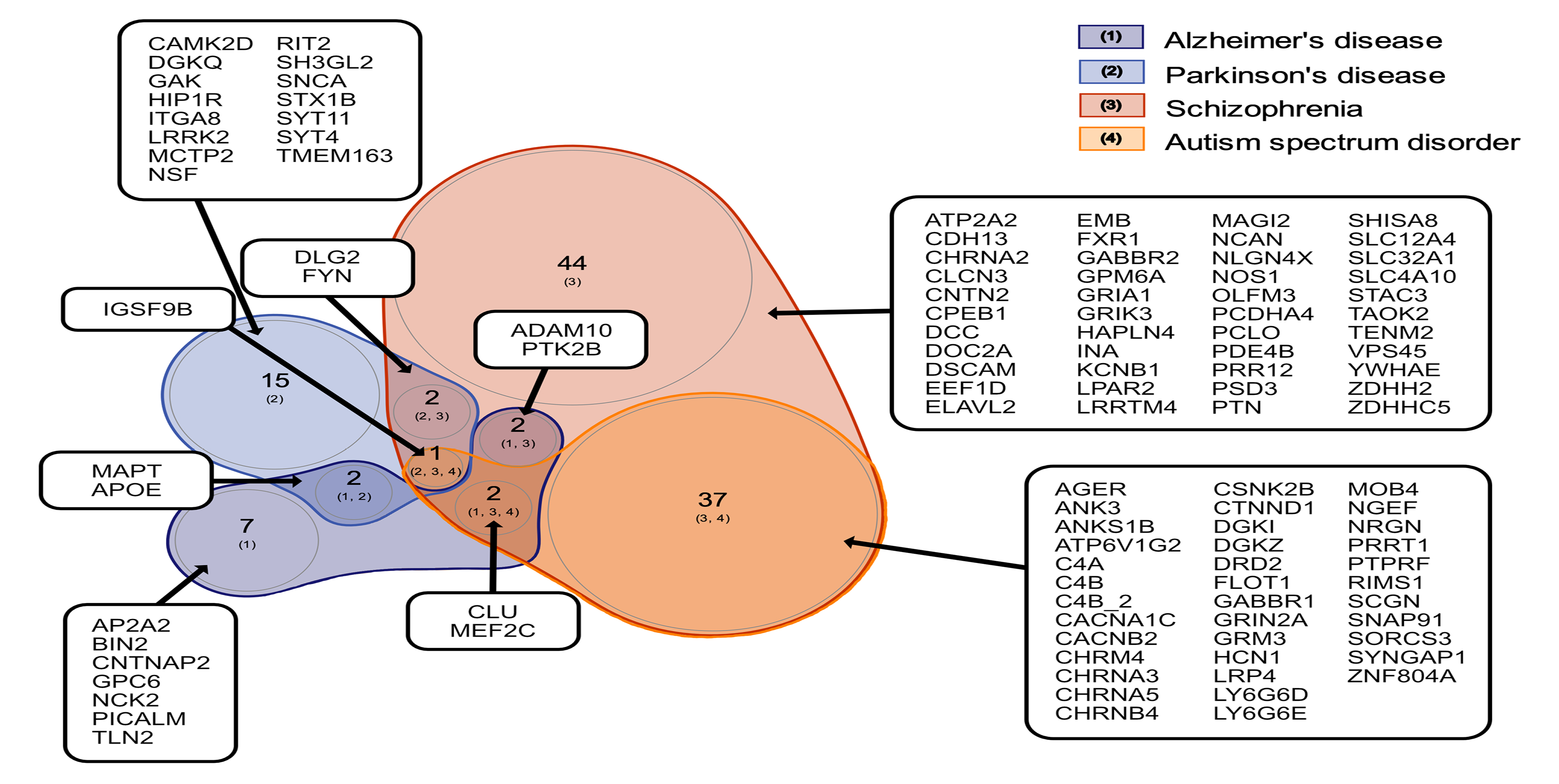
Overlap between genes associated with neurodevelopmental disorders and neurodegenerative diseases in humans based on genome-wide association study (GWAS) findings focusing on those overlapping with the GO synapse. Venn diagram visualizing overlap between synaptic suggestive candidate genes associated with the selected neurodevelopmental disorders and neurodegenerative diseases. The synaptic suggestive candidate genes, APOE and MAPT were associated with both Alzheimer’s and Parkinson’s disease, ADAM10 and PTK2B with both Alzheimer’s disease and schizophrenia, DLG2 and FYN with both Parkinson’s disease and schizophrenia, CLU and MEF2C with Alzheimer’s disease, schizophrenia and ASD, and IGSF9B with Parkinson’s disease, schizophrenia and ASD. See Table S2 for full lists of genes.

Comparison of the synaptic suggestive GWAS candidate genes across other combinations of diseases and disorders (Table S2) revealed the following overlapping findings: (i) apolipoprotein E (*APOE*) and microtubule associated protein tau (*MAPT*) for Parkinson’s disease and Alzheimer’s disease, (ii) ADAM metallopeptidase domain 10 (*ADAM10*) and protein tyrosine kinase 2 beta (*PTK2B*) for Alzheimer’s disease and schizophrenia, (iii) clusterin (*CLU*) and myocyte enhancer factor 2C (*MEF2C*) for Alzheimer’s disease, schizophrenia and ASD, (iv) discs large MAGUK scaffold protein 2 (*DLG2*) and FYN proto-oncogene, Src family tyrosine kinase (*FYN*) for Parkinson’s disease and schizophrenia, and (v) immunoglobulin superfamily member 9B (*IGSF9B*) for Parkinson’s disease, schizophrenia & ASD (Figures 2-3; Table S2). Based on the STRING database (v11.5), the majority of these genes are directly or indirectly connected either through PPIs (identified from experimental/biochemical data or reported in curated databases) or co-mentioning in PubMed abstracts, indicating involvement in similar biological mechanisms. PANX1 is connected to this STRING network via the proto-oncogene tyrosine-protein kinase (*Src*), which regulates GLUN2A and GLUN2B receptors and synaptic metaplasticity [69] and mediates PANX1-NMDA receptor crosstalk in pathophysiological synaptic plasticity [70,71] through physical interaction [70]. Given that PANX1 is connected to several of these neurological conditions and has been reported to be enriched at post-synaptic membranes, we next further characterized the neural PANX1 interactome (originally identified by our lab [11,13] by performing PANTHER pathway and PPI analyses to expand our understanding of its biological roles.

### Bioinformatics analyses of the PANX1 interactome revealed involvement of genes associated with cell structure regulation, proteostasis, neurodegeneration and synaptic enrichment

In addition to revealing possible biological roles of the PANX1 interactome, we also conducted pathway and PPI analyses to better understand (i) potential implications in disease and (ii) interactions between the PANX1-interacting proteins, based on existing knowledge. PANTHER enrichment analysis of the 483 *Mus musculus* PANX1-interacting proteins (unique UniProt accession numbers) revealed enrichment for six PANTHER pathways, namely the ubiquitin proteasome pathway (P00060, p-value = 3.65E-04), Parkinson’s disease (P00049, p-value = 1.05E-03), integrin signalling pathway (P00034, p-value = 1.61E-03), nicotinic acetylcholine receptor signaling pathway (P00044, p-value = 3.93E-03), inflammation mediated by chemokine and cytokine signaling pathway (P00031, p-value = 1.93E-02) and Huntington disease (P00029, p-value = 1.99E-02). An overview of the PANX1-interacting proteins annotated to the Parkinson’s disease PANTHER pathway can be found in Table 1.

**Table 1:** *Mus Musculus* PANX1-interacting proteins annotated to the Parkinson disease PANTHER pathway (P00049), other PANTHER pathways, and PANTHER protein classes.

| Gene symbol | Gene description | PANTHER protein class | PANTHER pathway |  |  |
| --- | --- | --- | --- | --- | --- |
|  |  |  | Apoptosis signaling pathway | FGF signaling pathway* | CCKR signaling map** |
| Hspa1b | Heat shock 70 kDa protein 1B | Hsp70 family chaperone | x |  |  |
| Hspa1l | Heat shock 70 kDa protein 1-like | Hsp70 family chaperone | x |  |  |
| Hspa2 | Heat shock-related 70 kDa protein 2 | Hsp70 family chaperone | x |  |  |
| Mapk1 | Mitogen-activated protein kinase 1 | Protein modifying enzyme | x | x | x |
| Mapk3 | Mitogen-activated protein kinase 3 | Protein modifying enzyme | x | x | x |
| Mcm5 | DNA replication licensing factor MCM5 | DNA metabolism protein | x |  |  |
| Psma3 | Proteasome subunit alpha type-3 | Protein modifying enzyme |  |  |  |
| Septin1 | Septin-1 | Cytoskeletal protein |  |  |  |
| Septin2 | Septin-2 | Cytoskeletal protein |  |  |  |
| Sfn | 14-3-3 protein sigma | Scaffold/adaptor protein |  | x |  |
| Ywhab | 14-3-3 protein beta/alpha | Scaffold/adaptor protein |  | x | x |

*Abbreviations:* CCKR, cholecystokinin receptor; FGF, fibroblast growth factor.
\* Involved in nervous system development and maintenance, neuroinflammation and dopaminergic neuron survival rate [112].
\*\* Patients with Parkinson's disease are more likely to hallucinate if having certain genetic polymorphisms in the CCK gene, and especially when combined with the CCKR, here CCKAR [113].

As PANX1 is involved in a wide range of synaptic functions, and synaptic dysfunction has been associated with neurological conditions such as Parkinson’s disease [72], we decided to identify synaptic PANX1-interacting proteins (∼18% of the neural PANX1 interactome) via cross-analysis with the *Mus musculus* cellular component GO term ‘synapse’. Using the 89 identified synaptic PANX1-interacting proteins as input, we created a STRING PPI network formed by 66 synaptic PANX1-interacting proteins (Figure 4) and identified 4 clusters involved in gene expression and translation, cytoskeleton organization, vesicle-mediated transport, and cell communication and its regulation, respectively (Table 2). When conducting the cross-analysis with both the *Mus musculus* and *Homo sapiens* synapse GO term (combined), 92 synaptic PANX1-interacting proteins were identified (see Table 3 for an overview of these proteins).

**Fig. 4:**
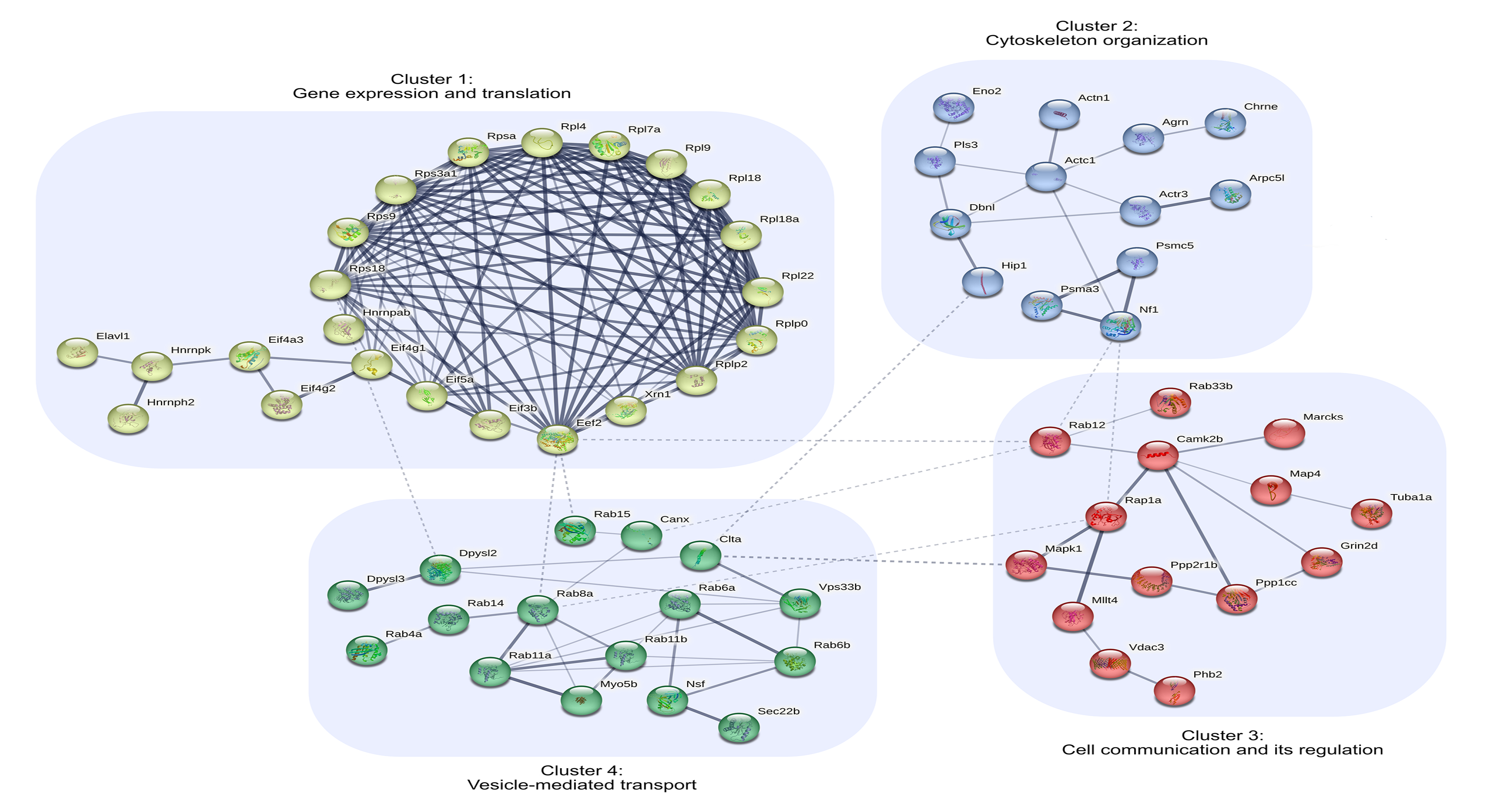
STRING Protein-Protein Interaction (PPI) network to explore the synaptic *Mus Musculus* PANX1 interactome and to identify clusters (potential functions presented in Table 2). A total of 66 synaptic PANX1-interacting proteins formed the PPI network (∼74% of the synaptic *Mus Musculus* PANX1 interactome), based on our chosen methodology. The thickness of the edges in the PPI network indicates the strength of data support and dashed line edges reveal PPIs between clusters.

**Table 2:** Synaptic Mus Musculus PANX1-interacting proteins forming the STRING Protein-Protein interaction network clusters annotated to selected PANTHER protein classes and Gene Ontology (GO) Biological Process (GP) terms.

| PANTHER protein class (GO BP term) | % of proteins |  |  |  |
| --- | --- | --- | --- | --- |
|  | Cluster 1 | Cluster 2 | Cluster 3 | Cluster 4 |
|  | n = 23 | n = 13 | n = 14 | n = 16 |
| Anatomical structure development (GO:0048856) | 26.1 | 53.8 | 71.4 | 37.5 |
| Nervous system development (GO:0007399) | 8.7 | 30.8 | 50.0 | 31.3 |
| Gene expression (GO:0010467) | 87.0 | 0.0 | 7.1 | 0.0 |
| Translational protein (PC00263) | 78.3 | 0.0 | 0.0 | 0.0 |
| Protein transport (GO:0015031) | 8.7 | 7.7 | 28.6 | 81.3 |
| Cytoskeleton organization (GO:0007010) | 0.0 | 61.5 | 21.4 | 25.0 |
| Cytoskeletal protein (PC00085) | 0.0 | 53.8 | 14.3 | 6.3 |
| Cell communication (GO:0007154) | 4.3 | 38.5 | 64.3 | 25.0 |
| Regulation of cell communication (GO:0010646) | 13.0 | 23.1 | 57.1 | 18.8 |
| Regulation of signaling (GO:0023051) | 13.0 | 23.1 | 57.1 | 25.0 |
| Vesicle-mediated transport (GO:0016192) | 0.0 | 15.4 | 28.6 | 93.8 |

**Table 3:**
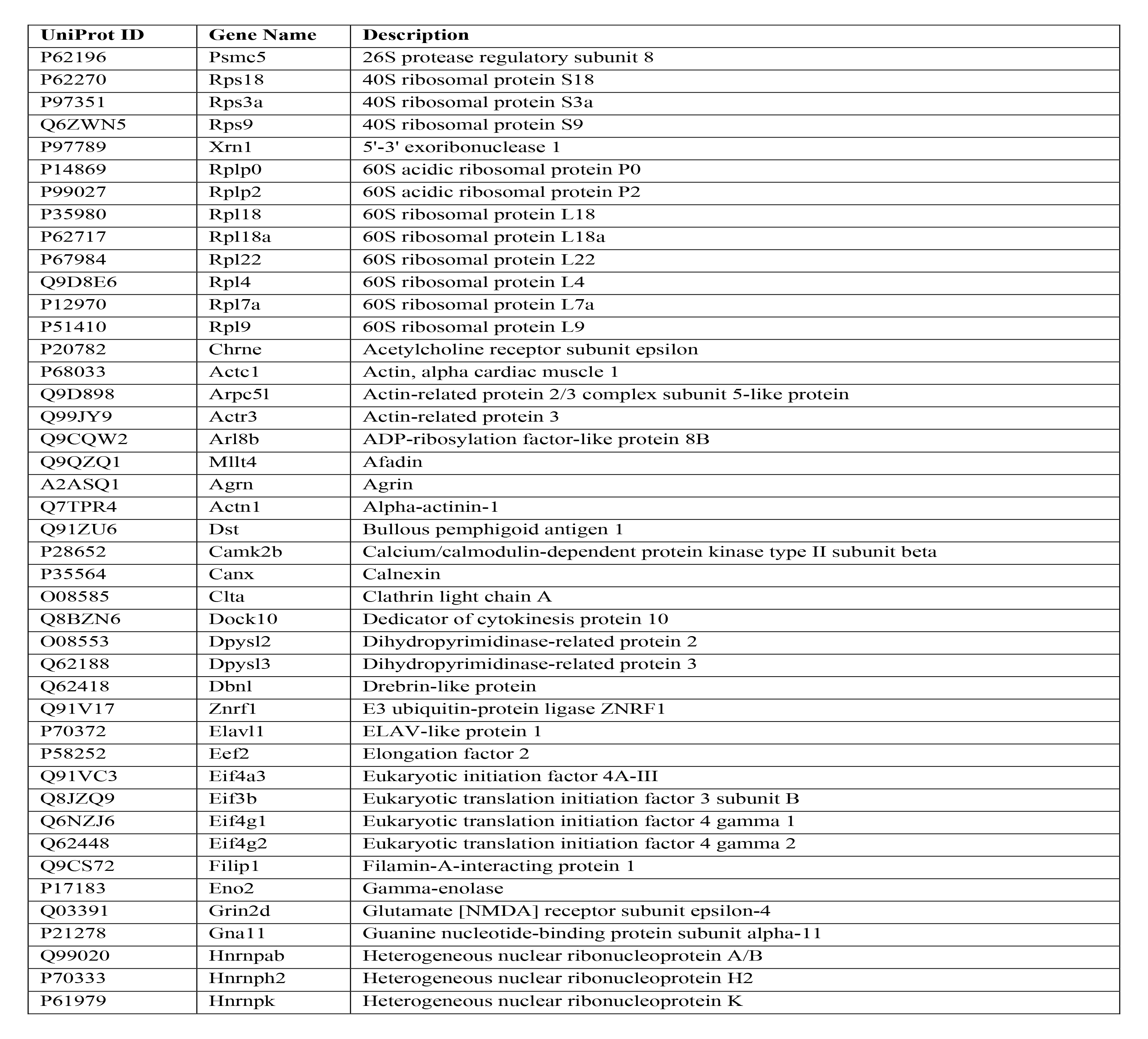
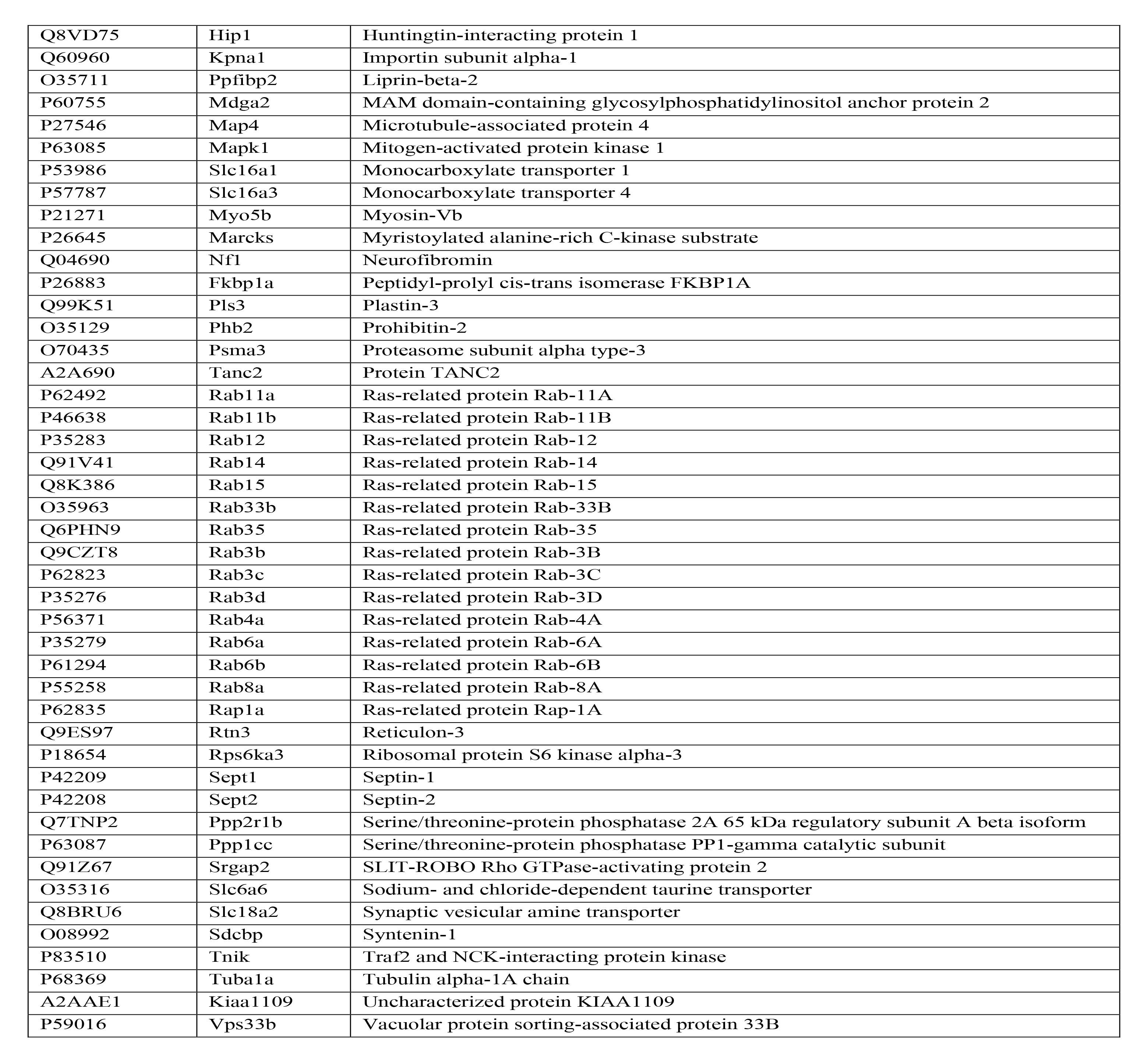
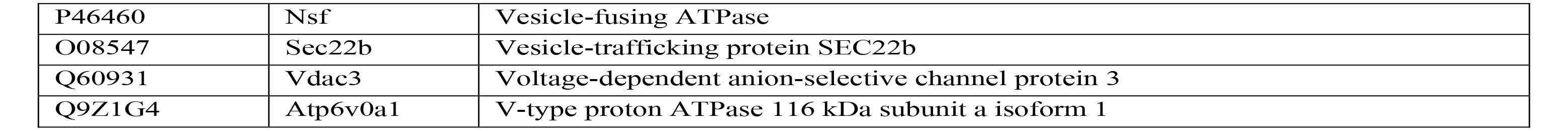
Overview of *Mus Musculus* synaptic PANX1-interacting proteins.

### Cross-analyses revealed overlap between the full neural and synapse-specific PANX1 interactome and neurological disease susceptibility genes

To explore connections between PANX1 and the four diseases of interest, we compared the entire PANX1 interactome with the suggestive candidate genes for each neurological disease (Figure 2). Overall, the two neurodevelopmental disease susceptibility gene sets exhibited a relatively larger degree of overlap with the PANX1 interactome than did the neurodegenerative disease susceptibility gene sets. Twenty schizophrenia susceptibility genes overlapped with the PANX1 interactome, which was the largest number in common, while ASD had 12 genes in common (see Table 4 for an overview). Several of these genes are involved in protein folding (GO:0006457; *Hspa1b, Hspa1l, Hspd1, Hspe1 and St13*) or regulation of translation (GO:0006417; Cnot1, *Ddx39b, Etf1, Lsm1 and Mapk3*).

**Table 4:** Genes encoding for PANX1-interacting proteins that have also been identified as suggestive GWAS candidate genes (Figure 2).

| Gene symbol | Gene description |
| --- | --- |
| **Parkinson's disease** |  |
| Nsf | Vesicle-fusing ATPase |
| **Alzheimer's disease** |  |
| Usp6nl | USP6 N-terminal-like protein |
| **Schizophrenia** |  |
| Acads* | Short-chain specific acyl-CoA dehydrogenase, mitochondrial |
| Actr1a* | Alpha-centractin |
| Agpat1* | 1-acyl-sn-glycerol-3-phosphate acyltransferase alpha |
| Ddx39b* | Spliceosome RNA helicase Ddx39b |
| Cnot1* | CCR4-NOT transcription complex subunit 1 |
| Csrp2 | Cysteine and glycine-rich protein 2 |
| Etf1 | Eukaryotic peptide chain release factor subunit 1 |
| Gmip | GEM-interacting protein |
| Gna12 | Guanine nucleotide-binding protein subunit alpha-12 |
| Gnl1* | Guanine nucleotide-binding protein-like 1 |
| Gulo | L-gulonolactone oxidase |
| Hspa1b* | Heat shock 70 kDa protein 1B |
| Hspa1l* | Heat shock 70 kDa protein 1-like |
| Hspd1* | 60 kDa heat shock protein, mitochondrial |
| Hspe1* | 10 kDa heat shock protein, mitochondrial |
| Lsm1 | U6 snRNA-associated Sm-like protein LSm1 |
| Mapk3 | Mitogen-activated protein kinase 3 |
| Plcl1* | Inactive phospholipase C-like protein 1 |
| Psmd6 | 26S proteasome non-ATPase regulatory subunit 6 |
| St13* | Hsc70-interacting protein |

\* Suggestive GWAS candidate genes associated with autism spectrum disorder that overlapped with the PANX1 interactome (complete overlap with the suggestive GWAS candidate genes associated with schizophrenia).

To identify synaptic-specific associations, we next compared the synaptic PANX1 interactome with synaptic suggestive GWAS candidate genes (Figure 3 and Table S2). Based on our methodological workflow (*e.g.,* restricted to genes reported more than once in the GWAS catalog and genes annotated to the *Mus Musculus* and *Homo Sapiens* synapse GO term), we identified one overlap: the gene coding for N-ethylmaleimide sensitive factor, vesicle-fusing ATPase (NSF) is both a suggestive candidate gene for Parkinson’s disease and a PANX1-interacting protein.

To explore possible synaptic PANX1-related disease connections more broadly, we next compared GO terms enriched within the synaptic PANX1 interactome with those enriched for the neurological conditions (based on synaptic disease susceptibility genes; Table 5). We included terms associated with the GO domains ‘cellular component’ and ‘biological process’ (choosing domains fitting the aim of this study), as these two relatively broader GO domains could help provide insight into common disease pathophysiology and potential PANX1 involvement. The ‘cellular component’ domain was selected to explore localization to structures associated with synapses, and/or structures influencing synapses and their development or stability (e.g., ‘somatodendritic tree’ should have some bearing on dendritic spines, ‘cytoplasmic vesicle’ would be associated with transport processes required to bring cargo to nascent spines, and so on). The following GO terms were enriched for the synaptic PANX1 interactome and each of the neurological conditions: Asymmetric synapse (GO:0032279), glutamatergic synapse (GO:0098978), neuron to neuron synapse (GO:0098984), postsynapse (GO:0098794), postsynaptic density (GO:0014069) and postsynaptic specialization (GO:0099572; Table 5).

**Table 5:**
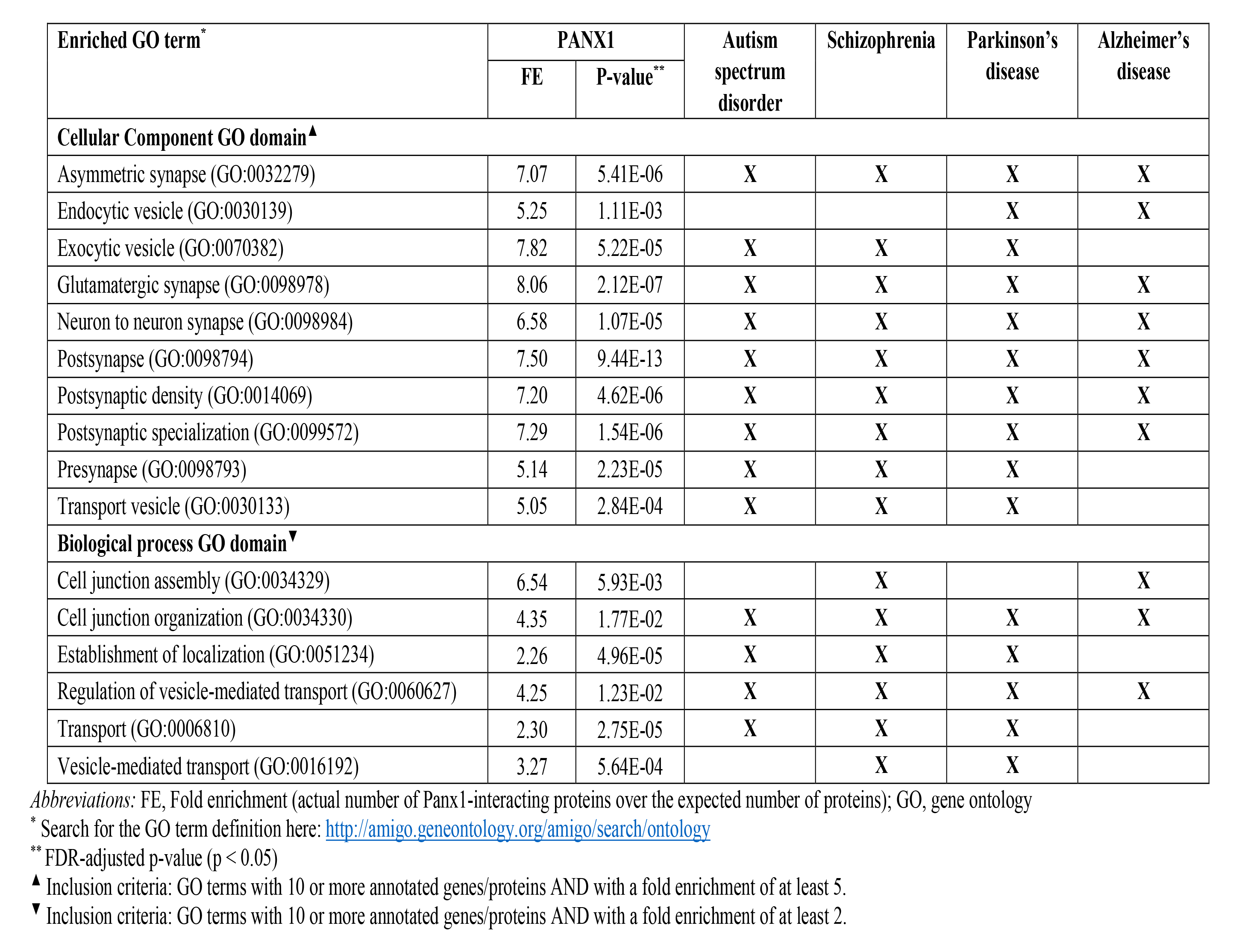
Enrichment analysis (GO terms within the ‘cellular component’ and ‘biological process’ GO domains) for the synaptic PANX1 interactome showing GO terms also found for the investigated neurological conditions (as indicated with X).

Focusing on the GO ‘biological process’, two GO terms overlapped (*i.e.,* cell junction organization (GO:0034330) and regulation of vesicle-mediated transport (GO:0060627)) when comparing the synaptic PANX1 interactome and the neurological conditions (Table 5).

### Specific human brain regions associated with ASD, schizophrenia, Parkinson’s disease, and Alzheimer’s disease

Since the neurological conditions we investigated are well-known to exhibit spatiotemporal specificity in terms of progression in affected brain regions (for example, Parkinson’s disease initially presents with striatal dysfunction), we next investigated regional specificity in elevated transcript expression for the synaptic PANX1 interactome and synaptic GWAS candidate genes by means of cross-analysis. Note that a gene was considered as “expressed” in a given brain region when the normalized transcript per million (nTPM; i.e., expression value) was above 1. Using the Human Protein Atlas v22.0 (<u>proteinatlas.org</u>) [66,67], we found three of the synaptic PANX1-interactors exhibited regionally-elevated transcript expression: RPL9 in cerebral cortex, SLC18A2 in pons, midbrain and hypothalamus, and TANC2 in cerebral cortex, hippocampal formation, and amygdala (Table 6; based on dataset on summarized expression in main brain regions not the specific expression in the more than 200 regions, areas and subfields, separately). Regionally elevated expression of synaptic neurological disease susceptibility genes was also found, with the highest number of genes noted for schizophrenia (16) and ASD (11; Table 6). Most of the disease-associated genes were elevated in the cerebral cortex, cerebellum, hippocampal formation and/or amygdala (Table 6).

**Table 6:** Overview of the overlap between the elevated genes in specific brain regions (Human Protein Atlas) and the synaptic PANX1 interactome/suggestive neurological disease susceptibility genes.

| Elevated gene | Brain region |  |  |  |  |  |  |  |
| --- | --- | --- | --- | --- | --- | --- | --- | --- |
|  | Cerebral cortex | Cerebellum | Hippocampal formation | Amygdala | Pons | Midbrain | Hypothalamus | Thalamus |
| CHRM4 | SCZ, ASD |  |  |  |  |  |  |  |
| CHRNA2 |  |  |  |  |  |  |  | SCZ |
| DGKZ | SCZ, ASD |  | SCZ, ASD | SCZ, ASD |  |  |  |  |
| GRIN2A | SCZ, ASD |  |  |  |  |  |  |  |
| HCN1 | SCZ, ASD |  |  |  |  |  |  |  |
| IGSF9B |  | PD, SCZ, ASD |  |  |  |  |  |  |
| KCNB1 | SCZ |  |  |  |  |  |  |  |
| MCTP2 | PD |  |  |  |  |  |  |  |
| MEF2C | AD, SCZ, ASD |  | AD, SCZ, ASD | AD, SCZ, ASD |  |  |  |  |
| NRGN | SCZ, ASD |  |  |  |  |  |  |  |
| PCLO |  | SCZ |  |  |  |  |  |  |
| PRRT1 | SCZ, ASD |  | SCZ, ASD | SCZ, ASD |  |  |  |  |
| RIMS1 |  | SCZ, ASD |  |  |  |  |  |  |
| RPL9 | PANX1 |  |  |  |  |  |  |  |
| SCGN |  | SCZ, ASD | SCZ, ASD |  |  |  | SCZ, ASD |  |
| SHISA8 |  | SCZ |  |  |  |  |  |  |
| SLC18A2 |  |  |  |  | PANX1 | PANX1 | PANX1 |  |
| SLC4A10 | SCZ |  |  |  |  |  |  |  |
| STX1B | PD |  |  |  |  |  |  |  |
| SYNGAP1 | SCZ, ASD |  | SCZ, ASD | SCZ, ASD |  |  |  |  |
| TANC2 | PANX1 |  | PANX1 | PANX1 |  |  |  |  |

## DISCUSSION

The goal of this study was to identify common synaptic genes and molecular pathways amongst neurological conditions and PANX1 using bioinformatics approaches. Using large-scale bioinformatics approaches have many advantages including but not limited to the: (i) ability to efficiently analyze a large amount of data, (ii) approaching analysis beyond tradition (e.g., data integration to get new insights), and (iii) possibility of making novel discoveries. Our study revealed multiple potential links between the PANX1 interactome and the investigated neurological conditions that now warrants validation. Overall, the two investigated neurodevelopmental disorders exhibited the largest overlap in synaptic susceptibility genes and were more abundantly represented in the PANX1 interactome. These results suggest that the molecular mechanisms underlying synaptic dysfunction in neurodevelopmental disorders may be more closely linked to one another and to PANX1 than is the case for neurodegenerative diseases, although it should be noted that this larger overlap also may be influenced by the size of these gene sets compared with those extracted for the investigated neurogenerative diseases.

### Further evidence for common genetic elements of ASD and schizophrenia

Notably, several of the susceptibility genes common to ASD and schizophrenia in our analysis have previously been linked to both disorders [73,74] and play key roles at the synapse (reviewed in [1,3,4]). For example, ANK3 (ANKG protein), GRIN2A (GluN2A), and SYNGAP1 have established synaptic roles [75–78] and connections to ASD, schizophrenia, and other co-morbid neurodevelopmental disorders like epilepsy [79–84]. These three genes were not identified in our PANX1 interactome; however, we identified another GluN-coding gene, *GRIN2D*. Given PANX1 has been previously reported to form a complex with GluN1 (obligatory NMDA receptor subunit) and Src [70], the relatively immature state of the Neuro2a cell model could account for the lack of identification of GluN2a as a PANX1 interactor in our system. Conversely, the lipid raft scaffolding protein, flotillin was common to ASD and schizophrenia, but has not been extensively studied in either neurodevelopmental disorder. Given lipid raft/cholesterol dysfunction has been implicated in the monogenic common ASD Fragile X syndrome and has been linked to glutamatergic synapse formation [85], further exploration of the role of flotillin in synaptic dysfunction in ASD and schizophrenia could be informative. Further analysis is now warranted to explore the shared synaptic genetic susceptibility between ASD and schizophrenia, including individual variability in risk, onset, severity and progression. Some studies have looked into this shared genetic basis. For instance, significant genetic correlations have been identified between ASD and schizophrenia (but not between Alzheimer’s disease and Parkinson’s disease)[68], similar to what we observed in this study. Even though ASD and schizophrenia share genetic risks, they seem to have distinct developmental profiles [86]; multi-omics studies could help tease out the underpinnings.

The established synaptic roles of these neurological disease susceptibility genes and our recent discovery of PANX1 regulation of dendritic spine stability prompted our synaptic PANX1 interactome STRING analysis, which identified clusters in gene expression and translation, cytoskeleton organization, vesicle-mediated transport, and cell communication and its regulation (Table 2). Dysregulation in these same cellular processes are observed in ASD [87,88], schizophrenia [89,90], Parkinson’s disease [91,92], and Alzheimer’s disease [93,94], and PANX1 has been linked to these conditions (reviewed in [95,96]). Of particular note, there are >40 non-coding *PANX1* variants in the VariCarta ASD database (https://varicarta.msl.ubc.ca/index [97]) and there are several ASD-linked *PANX1* SNPs associated with brain-specific *PANX1* gene expression changes [98]. Our cross-analysis of PANX1 interactors, synaptic-expressed genes, and genetic risk susceptibility to neurological conditions is a key step in bridging our gaps in understanding how PANX1 is linked to these various conditions.

Given the key synaptic findings from our broader analyses, we thought it prudent to refine our approach with the GO synaptic gene set; however, the results were somewhat surprising. Despite the strong representation of well-established and important synaptic genes associated with neurological conditions in the PANX1 set (refer to Table 3, e.g., *Dpysl2*, *Actr3*, *Hip1*, several *Rab*s etc.), when we restricted our disease-associated analysis to ‘GO synaptic’ genes, we only identified the vesicular trafficking regulator NSF as the sole common gene shared by PANX1 and a neurological disease susceptibility gene set (specifically for Parkinson’s disease). Given that studies in Drosophila have shown that expression of a dominant-negative form of Nsf2 leads to disrupted neuromuscular junction and synaptic structural development [99,100] and that this is linked to actin-cytoskeleton regulation [101], this finding suggests that investigating the NSF-PANX1 interaction could be of key interest in terms of exploring a putative role for PANX1 in synaptic dysfunction in Parkinson’s disease [102]. The connection to Parkinson’s disease is further bolstered by our PANTHER pathway analysis, which identified Parkinson’s disease as one of six pathways exhibiting enrichment within the PANX1 interactome. Importantly, the limited output of the GO synaptic analysis (1 gene identified) underscores a key caveat of bioinformatics and GO, in that many key connections may be missed as the outputs are limited by the inputs (a gene may have not met overly restrictive inclusion criteria or certain literature has been missed). Given this limitation, we broadened our approach to compare enriched synapse-specific GO terms (Table 5). In accordance with casting a wide net, we focused on cellular component locations (‘cellular component’ domain) and biological processes (‘biological process’ domain), which are relatively broader than the third GO domain, ‘molecular function’. We identified similar cellular components/processes as those identified in our STRING analysis of PANX1 PPIs, such as vesicle-mediated transport. These similarities provide potential avenues of insight into the connection between PANX1 and various neurological conditions, as discussed above. Further dissection of the brain region-specific could also provide important clues.

### Limitations of GWAS and GO analysis can obscure key synaptic PPIs

GWAS exhibit several key caveats. Importantly, GWAS were historically somewhat restricted to European populations, thereby limiting their broader use and application [103]; importantly, some recent progress has been made in expanding GWAS to more groups, especially Asian populations (see Table 7 for details of GWASs used in the current study). Additionally, although GWAS identify genetic differences associated with disease, these are not necessarily causal, and where mechanistic links might be present, these can be challenging and complicated to unravel [104]. In terms of the analysis of synaptic representation within suggestive candidate genes for neurological conditions, there are several types of connections that can be missed, as briefly discussed above. For example, given the limitations of gene ontologies [105], not all proteins that could be present at synapses during their lifespan may have their genes annotated to GO synapse – for example *PANX1* itself is not found in GO synapse, despite its described localization and functional characterization at the post-synapse. Similarly, a recently created synaptic gene ontology database (https://www.syngoportal.org/) does not contain *PANX1*. Furthermore, identification of a connection does not imply causality; this requires careful and in-depth follow up cell biology studies amongst other analyses. Additionally, the analysis is limited to gene level associations with disease and does not account for disease-associated differences in expression levels, post-translational modifications or protein-targeting pathophysiological mechanisms, like autoantibody production, all of which have been described for CRMP2 (e.g., changes in protein expression levels in several neurological conditions [106], hyperphosphorylation in Alzheimer’s disease [107–109], and auto-antibodies in autism spectrum disorders [110]). Finally, these genetic approaches cannot account for the influence of environmental factors such as inflammation due to injury or infection, which would be expected to have a major impact on PANX1 function and regulation [95,111].

**Table 7:** Overview of the specified study populations in the Genome-Wide Association Studies (GWASs) extracted from the GWAS catalog for each neurological condition (prior to applying filtering criteria).

| **Populations** | **Autism spectrum disorder (n = 49)** | **Schizophrenia (n = 212)** | **Parkinson's disease (n = 69)** | **Alzheimer's disease (n = 161)** |
| --- | --- | --- | --- | --- |
| European | 40.8% | 45.3% | 73.9% | 59.0% |
| East Asian | 8.2% | 19.8% | 8.7% | 6.2% |
| South Asian | - | 0.9% | 1.4% | - |
| Asian unspecified | 2.0% | 2.8% | 4.3% | - |
| Greater Middle Eastern (Middle Eastern, North African or Persian) | 4.1% | 0.5% | - | 2.5% |
| African American or Afro-Caribbean | 2.0% | 5.7% | - | 6.2% |
| Hispanic or Latin American | - | 2.8% | 2.9% | 2.5% |
| Native American | - | 0.9% | - | 0.6% |
| Sub-Saharan African | - | 0.5% | 1.4% | - |
| African unspecified | 2.0% | 1.4% | - | 0.6% |
| Oceanian | - | 0.9% | - | - |
| Other | 2.0% | 2.4% | 2.9% | 0.6% |

### Where do we go from here?

To fully appreciate both the complexity and validity of putative pathophysiological mechanisms hinted at by the analyses presented here, additional work in rodent and human iPSC-derived models of neuropsychiatric disease would be a logical next step. For example, while we have previously validated interactions of PANX1 with ARP3 and CRMP2 in cell lines, investigating their interplay within living neurons and brain will help to shed light on how this and related PPI networks intersect in synapse development in health and disease states.

## ACKNOWLEDGMENTS

This project was supported by operating grants from the Canadian Institutes of Health Research (MOP142215), the Natural Sciences and Engineering Research Council of Canada (NSERC; 402270- 2011), and the University of Victoria Division of Medical Sciences to LAS. LAS was also supported by a Michael Smith Foundation for Health Research and British Columbia Schizophrenia Society Foundation Scholar Award (5900). LEWS was supported by a Vanier Canada Graduate Scholarship (NSERC). Additionally, the authors are grateful Juan C. Sanchez-Arias for his initial input on the study design and discussion for the first bioRxiv preprint version of this work (posted October 11, 2019). LAS and SDF designed the study with inputs from LEWS. LEWS identified the PANX1 interactome in mouse N2a cells, SDF performed the bioinformatics analyses, and SDF, LEWS and LAS wrote the manuscript. All authors approved the final manuscript.

## DECLARATION OF INTERESTS

The authors have no conflicts of interest to declare, and the study itself has been exempted from ethical review by the University of Victoria’s Human Research Ethics Board.

## Notes

### Competing Interest Statement

The authors have declared no competing interest.

### Summary of Updates

Added key missing reference to Leigh Wicki-Stordeur's PhD thesis.

