## Supplemental Tables for "Overlap in synaptic neurological condition susceptibility pathways and the neural pannexin 1 interactome revealed by bioinformatics analyses"

**Table S1: Overview of the Genome-Wide Association Studies (GWAS) extracted from the GWAS catalog by means of the R package gwasrapidd**

| disease | pubmed_id | author_fullname | publication_date | title | publication |
|---|---|---|---|---|---|
| Austism spectrum disorder | 23453885 | Smoller JW | 02-27-2013 | Identification of risk loci with shared effects on five major psychiatric disorders: a genome-wide analysi | Lancet |
| Austism spectrum disorder | 24574247 | Yuan H | 02-25-2014 | Investigation of maternal genotype effects in autism by genome-wide association. | Autism Res |
| Austism spectrum disorder | 21519539 | Cho SC | 02-25-2011 | Genome-wide association scan of korean autism spectrum disorders with language delay: a preliminary | Psychiatry Investig |
| Austism spectrum disorder | 25534755 | Chaste P | 09-30-2014 | A genome-wide association study of autism using the Simons Simplex Collection: Does reducing phenot | Biol Psychiatry |
| Austism spectrum disorder | 26398136 | Kuo PH | 09-23-2015 | Genome-Wide Association Study for Autism Spectrum Disorder in Taiwanese Han Population. | PLoS One |
| Austism spectrum disorder | 26314684 | Liu X | 08-28-2015 | Genome-wide Association Study of Autism Spectrum Disorder in the East Asian Populations. | Autism Res |
| Austism spectrum disorder | 28540026 | Anney RJL | 05-22-2017 | Meta-analysis of GWAS of over 16,000 individuals with autism spectrum disorder highlights a novel loc | Mol Autism |
| Austism spectrum disorder | 28641744 | Guo W | 07-01-2017 | Polygenic risk score and heritability estimates reveals a genetic relationship between ASD and OCD. | Eur Neuropsychopharm |
| Austism spectrum disorder | 30804558 | Grove J | 02-25-2019 | Identification of common genetic risk variants for autism spectrum disorder. | Nat Genet |
| Austism spectrum disorder | 30692689 | Schork AJ | 01-28-2019 | A genome-wide association study of shared risk across psychiatric disorders implicates gene regulation | Nat Neurosci |
| Austism spectrum disorder | 31209380 | Demontis D | 06-17-2019 | Genome-wide association study implicates CHRNA2 in cannabis use disorder. | Nat Neurosci |
| Austism spectrum disorder | 30289880 | Qi G | 10-05-2018 | Heritability informed power optimization (HIPO) leads to enhanced detection of genetic associations ac | PLoS Genet |
| Austism spectrum disorder | 32606422 | Wu Y | 06-30-2020 | Multi-trait analysis for genome-wide association study of five psychiatric disorders. | Transl Psychiatry |
| Austism spectrum disorder | 31835028 | Cross-Disorder Grou | 12-01-2019 | Genomic Relationships, Novel Loci, and Pleiotropic Mechanisms across Eight Psychiatric Disorders. | Cell |
| Austism spectrum disorder | 32747698 | Matoba N | 08-03-2020 | Common genetic risk variants identified in the SPARK cohort support DDHD2 as a candidate risk gene fo | Transl Psychiatry |
| Austism spectrum disorder | 33479212 | Yao X | 01-21-2021 | Integrative analysis of genome-wide association studies identifies novel loci associated with neuropsycl | Transl Psychiatry |
| Austism spectrum disorder | 33686288 | Peyrot WJ | 03-08-2021 | Identifying loci with different allele frequencies among cases of eight psychiatric disorders using CC-GV | Nat Genet |
| Austism spectrum disorder | 34069769 | Al-Sarraj Y | 05-18-2021 | Family-Based Genome-Wide Association Study of Autism Spectrum Disorder in Middle Eastern Familie | Genes (Basel) |
| Austism spectrum disorder | 35215271 | Almandil NB | 01-27-2022 | Integration of Transcriptome and Exome Genotyping Identifies Significant Variants with Autism Spectru | Pharmaceuticals (Basel |
| Austism spectrum disorder | 35764056 | Rao S | 06-28-2022 | Genetic Relationships between Attention-Deficit/Hyperactivity Disorder, Autism Spectrum Disorder, and | Neuropsychobiology |
| Austism spectrum disorder | 35717853 | Baranova A | 06-13-2022 | Shared genetics between autism spectrum disorder and attention-deficit/hyperactivity disorder and the | Psychiatry Res |
| Schizophrenia | 25056061 | Ripke S | 07-22-2014 | Biological insights from 108 schizophrenia-associated genetic loci. | Nature |
| Schizophrenia | 22688191 | Bergen SE | 06-12-2012 | Genome-wide association study in a Swedish population yields support for greater CNV and MHC involv | Mol Psychiatry |
| Schizophrenia | 18282107 | Shifman S | 02-15-2008 | Genome-wide association identifies a common variant in the reelin gene that increases the risk of schiz | PLoS Genet |
| Schizophrenia | 17522711 | Lencz T | 03-20-2007 | Converging evidence for a pseudoautosomal cytokine receptor gene locus in schizophrenia. | Mol Psychiatry |
| Schizophrenia | 18677311 | O'Donovan MC | 07-30-2008 | Identification of loci associated with schizophrenia by genome-wide association and follow-up. | Nat Genet |
| Schizophrenia | 19197363 | Need AC | 02-06-2009 | A genome-wide investigation of SNPs and CNVs in schizophrenia. | PLoS Genet |
| Schizophrenia | 19571809 | Shi J | 07-01-2009 | Common variants on chromosome 6p22.1 are associated with schizophrenia. | Nature |
| Schizophrenia | 19571811 | Purcell SM | 07-01-2009 | Common polygenic variation contributes to risk of schizophrenia and bipolar disorder. | Nature |
| Schizophrenia | 19571808 | Stefansson H | 07-01-2009 | Common variants conferring risk of schizophrenia. | Nature |
| Schizophrenia | 20832056 | Ikeda M | 09-08-2010 | Genome-wide association study of schizophrenia in a Japanese population. | Biol Psychiatry |
| Schizophrenia | 20558996 | Ott J | 06-17-2010 | Pilot study on schizophrenia in Sardinia. | Hum Hered |
| Schizophrenia | 21679298 | Ma X | 06-16-2011 | A genome-wide association study for quantitative traits in schizophrenia in China. | Genes Brain Behav |
| Schizophrenia | 21747397 | Rietschel M | 07-12-2011 | Association between genetic variation in a region on chromosome 11 and schizophrenia in large sample | Mol Psychiatry |
| Schizophrenia | 21752600 | Chen J | 07-12-2011 | Two non-synonymous markers in PTPN21, identified by genome-wide association study data-mining and | Schizophr Res |
| Schizophrenia | 22037555 | Shi Y | 10-30-2011 | Common variants on 8p12 and 1q24.2 confer risk of schizophrenia. | Nat Genet |
| Schizophrenia | 23358160 | Børglum AD | 01-29-2013 | Genome-wide study of association and interaction with maternal cytomegalovirus infection suggests ne | Mol Psychiatry |
| Schizophrenia | 23894747 | Aberg KA | 02-01-2013 | A comprehensive family-based replication study of schizophrenia genes. | JAMA Psychiatry |
| Schizophrenia | 23453885 | Smoller JW | 02-27-2013 | Identification of risk loci with shared effects on five major psychiatric disorders: a genome-wide analysi | Lancet |
| Schizophrenia | 23594818 | Ikeda M | 04-17-2013 | Evidence for shared genetic risk between methamphetamine-induced psychosis and schizophrenia. | Neuropsychopharmacol |
| Schizophrenia | 22883433 | Irish Schizophrenia G | 08-07-2012 | Genome-wide association study implicates HLA-C*01:02 as a risk factor at the major histocompatibility | Biol Psychiatry |
| Schizophrenia | 23974872 | Ripke S | 08-25-2013 | Genome-wide association analysis identifies 13 new risk loci for schizophrenia. | Nat Genet |
| Schizophrenia | 22037552 | Yue WH | 10-30-2011 | Genome-wide association study identifies a susceptibility locus for schizophrenia in Han Chinese at 11p | Nat Genet |

| | | | | | |
|---|---|---|---|---|---|
| Schizophrenia | 23142968 | Betcheva ET | 11-07-2012 | Whole-genome-wide association study in the Bulgarian population reveals HHAT as schizophrenia susce | Psychiatr Genet |
| Schizophrenia | 21682944 | Alkelai A | 06-20-2011 | DOCK4 and CEACAM21 as novel schizophrenia candidate genes in the Jewish population. | Int J Neuropsychopharm |
| Schizophrenia | 21926974 | Ripke S | 09-18-2011 | Genome-wide association study identifies five new schizophrenia loci. | Nat Genet |
| Schizophrenia | 24166486 | Sleiman P | 10-29-2013 | GWAS meta analysis identifies TSNARE1 as a novel Schizophrenia / Bipolar susceptibility locus. | Sci Rep |
| Schizophrenia | 18347602 | Sullivan PF | 03-18-2008 | Genomewide association for schizophrenia in the CATIE study: results of stage 1. | Mol Psychiatry |
| Schizophrenia | 24253340 | Lencz T | 11-19-2013 | Genome-wide association study implicates NDST3 in schizophrenia and bipolar disorder. | Nat Commun |
| Schizophrenia | 24280982 | Ruderfer DM | 11-26-2013 | Polygenic dissection of diagnosis and clinical dimensions of bipolar disorder and schizophrenia. | Mol Psychiatry |
| Schizophrenia | 23212062 | Fanous AH | 12-01-2012 | Genome-wide association study of clinical dimensions of schizophrenia: polygenic effect on disorganize | Am J Psychiatry |
| Schizophrenia | 24043878 | Wong EH | 09-16-2013 | Common variants on Xq28 conferring risk of schizophrenia in Han Chinese. | Schizophr Bull |
| Schizophrenia | 24086445 | Wang Q | 09-24-2013 | Genome-wide association analysis with gray matter volume as a quantitative phenotype in first-episod | PLoS One |
| Schizophrenia | 26198764 | Goes FS | 07-21-2015 | Genome-wide association study of schizophrenia in Ashkenazi Jews. | Am J Med Genet B Neu |
| Schizophrenia | 25781172 | Avramopoulos D | 03-17-2015 | Infection and inflammation in schizophrenia and bipolar disorder: a genome wide study for interactions | PLoS One |
| Schizophrenia | 26531332 | Kim LH | 11-14-2015 | Genome-wide association study with the risk of schizophrenia in a Korean population. | Am J Med Genet B Neu |
| Schizophrenia | 28322246 | McLaughlin RL | 03-21-2017 | Genetic correlation between amyotrophic lateral sclerosis and schizophrenia. | Nat Commun |
| Schizophrenia | 28540026 | Anney RJL | 05-22-2017 | Meta-analysis of GWAS of over 16,000 individuals with autism spectrum disorder highlights a novel loc | Mol Autism |
| Schizophrenia | 28746715 | Smeland OB | 07-26-2017 | Identification of Genetic Loci Jointly Influencing Schizophrenia Risk and the Cognitive Traits of Verbal-N | JAMA Psychiatry |
| Schizophrenia | 28991256 | Li Z | 10-09-2017 | Genome-wide association analysis identifies 30 new susceptibility loci for schizophrenia. | Nat Genet |
| Schizophrenia | 26249676 | Wolthusen RP | 08-07-2015 | Genetic underpinnings of left superior temporal gyrus thickness in patients with schizophrenia. | World J Biol Psychiatry |
| Schizophrenia | 27922604 | Yu H | 12-06-2016 | Common variants on 2p16.1, 6p22.1 and 10q24.32 are associated with schizophrenia in Han Chinese po | Mol Psychiatry |
| Schizophrenia | 28744025 | Woolston AL | 07-25-2017 | Genetic loci associated with an earlier age at onset in multiplex schizophrenia. | Sci Rep |
| Schizophrenia | 30285260 | Ikeda M | 10-03-2018 | Genome-Wide Association Study Detected Novel Susceptibility Genes for Schizophrenia and Shared Tra | Schizophr Bull |
| Schizophrenia | 30470734 | Hackinger S | 11-23-2018 | Evidence for genetic contribution to the increased risk of type 2 diabetes in schizophrenia. | Transl Psychiatry |
| Schizophrenia | 31209380 | Demontis D | 06-17-2019 | Genome-wide association study implicates CHRNA2 in cannabis use disorder. | Nat Neurosci |
| Schizophrenia | 30610197 | Smeland OB | 01-04-2019 | Genome-wide analysis reveals extensive genetic overlap between schizophrenia, bipolar disorder, and i | Mol Psychiatry |
| Schizophrenia | 30920090 | Trochet H | 03-28-2019 | Bayesian meta-analysis across genome-wide association studies of diverse phenotypes. | Genet Epidemiol |
| Schizophrenia | 30692689 | Schork AJ | 01-28-2019 | A genome-wide association study of shared risk across psychiatric disorders implicates gene regulation | Nat Neurosci |
| Schizophrenia | 29330379 | Zuber V | 01-12-2018 | Identification of shared genetic variants between schizophrenia and lung cancer. | Sci Rep |
| Schizophrenia | 30626913 | Amare AT | 01-09-2019 | Bivariate genome-wide association analyses of the broad depression phenotype combined with major c | Mol Psychiatry |
| Schizophrenia | 31268507 | Periyasamy S | 07-03-2019 | Association of Schizophrenia Risk With Disordered Niacin Metabolism in an Indian Genome-wide Assoc | JAMA Psychiatry |
| Schizophrenia | 29121268 | Amare AT | 11-09-2017 | Association of Polygenic Score for Schizophrenia and HLA Antigen and Inflammation Genes With Respo | JAMA Psychiatry |
| Schizophrenia | 30289880 | Qi G | 10-05-2018 | Heritability informed power optimization (HIPO) leads to enhanced detection of genetic associations ac | PLoS Genet |
| Schizophrenia | 29483656 | Pardiñas AF | 02-26-2018 | Common schizophrenia alleles are enriched in mutation-intolerant genes and in regions under strong ba | Nat Genet |
| Schizophrenia | 30804558 | Grove J | 02-25-2019 | Identification of common genetic risk variants for autism spectrum disorder. | Nat Genet |
| Schizophrenia | 31155012 | Nakahara S | 06-03-2019 | Dentate gyrus volume deficit in schizophrenia. | Psychol Med |
| Schizophrenia | 31374203 | Lam M | 08-01-2019 | Pleiotropic Meta-Analysis of Cognition, Education, and Schizophrenia Differentiates Roles of Early Neur | Am J Hum Genet |
| Schizophrenia | 31579629 | Fiorica PN | 09-26-2019 | Transcriptome association studies of neuropsychiatric traits in African Americans implicate PRMT7 in s | PeerJ |
| Schizophrenia | 31591465 | Bigdeli TB | 10-07-2019 | Contributions of common genetic variants to risk of schizophrenia among individuals of African and Lati | Mol Psychiatry |
| Schizophrenia | 32762793 | Jang SK | 08-07-2020 | Genetic correlation, pleiotropy, and causal associations between substance use and psychiatric disorder | Psychol Med |
| Schizophrenia | 32606422 | Wu Y | 06-30-2020 | Multi-trait analysis for genome-wide association study of five psychiatric disorders. | Transl Psychiatry |
| Schizophrenia | 31740837 | Lam M | 11-18-2019 | Comparative genetic architectures of schizophrenia in East Asian and European populations. | Nat Genet |
| Schizophrenia | 31835028 | Cross-Disorder Grou | 12-01-2019 | Genomic Relationships, Novel Loci, and Pleiotropic Mechanisms across Eight Psychiatric Disorders. | Cell |
| Schizophrenia | 31735061 | Liu X | 11-16-2019 | Wnt receptor gene FZD1 was associated with schizophrenia in genome-wide SNP analysis of the Austra | Aust N Z J Psychiatry |
| Schizophrenia | 32589924 | Bi W | 06-23-2020 | A Fast and Accurate Method for Genome-Wide Time-to-Event Data Analysis and Its Application to UK B | Am J Hum Genet |
| Schizophrenia | 31913414 | Bahrami S | 01-08-2020 | Shared Genetic Loci Between Body Mass Index and Major Psychiatric Disorders: A Genome-wide Associ | JAMA Psychiatry |
| Schizophrenia | 32107650 | Liu L | 02-27-2020 | A trans-ethnic two-stage polygenetic scoring analysis detects genetic correlation between osteoporosis | Clin Transl Med |

| | | | | | |
|---|---|---|---|---|---|
| Schizophrenia | 32201043 | Smeland OB | 02-08-2020 | Genome-wide Association Analysis of Parkinson's Disease and Schizophrenia Reveals Shared Genetic Ar | Biol Psychiatry |
| Schizophrenia | 33479212 | Yao X | 01-21-2021 | Integrative analysis of genome-wide association studies identifies novel loci associated with neuropsycl | Transl Psychiatry |
| Schizophrenia | 32772156 | Muntané G | 08-09-2020 | The shared genetic architecture of schizophrenia, bipolar disorder and lifespan. | Hum Genet |
| Schizophrenia | 33686288 | Peyrot WJ | 03-08-2021 | Identifying loci with different allele frequencies among cases of eight psychiatric disorders using CC-GV | Nat Genet |
| Schizophrenia | 32965087 | Zamanpoor M | 09-23-2020 | The genetic basis for the inverse relationship between rheumatoid arthritis and schizophrenia. | Mol Genet Genomic Me |
| Schizophrenia | 33169155 | Bigdeli TB | 11-10-2020 | Genome-Wide Association Studies of Schizophrenia and Bipolar Disorder in a Diverse Cohort of US Vete | Schizophr Bull |
| Schizophrenia | 34099189 | Blokland GAM | 03-23-2021 | Sex-Dependent Shared and Nonshared Genetic Architecture Across Mood and Psychotic Disorders. | Biol Psychiatry |
| Schizophrenia | 33907183 | Guo S | 04-27-2021 | Genome wide association study identifies four loci for early onset schizophrenia. | Transl Psychiatry |
| Schizophrenia | 33133133 | Lee KY | 08-28-2020 | Genome-Wide Search for SNP Interactions in GWAS Data: Algorithm, Feasibility, Replication Using Schi | Front Genet |
| Schizophrenia | 34380480 | Liu J | 08-12-2021 | Genome-wide association study followed by trans-ancestry meta-analysis identify 17 new risk loci for s | BMC Med |
| Schizophrenia | 34159505 | Wang H | 06-16-2021 | Novel loci and potential mechanisms of major depressive disorder, bipolar disorder, and schizophrenia. | Sci China Life Sci |
| Schizophrenia | 35235886 | Song W | 02-17-2022 | Genome-wide identification of the shared genetic basis of cannabis and cigarette smoking and schizopl | Psychiatry Res |
| Schizophrenia | 34737426 | Jiang L | 11-04-2021 | A generalized linear mixed model association tool for biobank-scale data. | Nat Genet |
| Schizophrenia | 34594039 | Sakaue S | 09-30-2021 | A cross-population atlas of genetic associations for 220 human phenotypes. | Nat Genet |
| Schizophrenia | 35019943 | Pardiñas AF | 01-12-2022 | Interaction Testing and Polygenic Risk Scoring to Estimate the Association of Common Genetic Variants | JAMA Psychiatry |
| Schizophrenia | 34976021 | Li W | 12-17-2021 | Identification of a Risk Locus at 7p22.3 for Schizophrenia and Bipolar Disorder in East Asian Populations | Front Genet |
| Schizophrenia | 34662886 | Backman JD | 10-18-2021 | Exome sequencing and analysis of 454,787 UK Biobank participants. | Nature |
| Schizophrenia | 35396580 | Trubetskoy V | 04-08-2022 | Mapping genomic loci implicates genes and synaptic biology in schizophrenia. | Nature |
| Parkinson disease | 17052657 | Fung HC | 09-28-2006 | Genome-wide genotyping in Parkinson's disease and neurologically normal controls: first stage analysis | Lancet Neurol |
| Parkinson disease | 18985386 | Pankratz N | 11-06-2008 | Genomewide association study for susceptibility genes contributing to familial Parkinson disease. | Hum Genet |
| Parkinson disease | 19915575 | Simón-Sánchez J | 11-15-2009 | Genome-wide association study reveals genetic risk underlying Parkinson's disease. | Nat Genet |
| Parkinson disease | 20070850 | Edwards TL | 01-13-2010 | Genome-wide association study confirms SNPs in SNCA and the MAPT region as common risk factors fc | Ann Hum Genet |
| Parkinson disease | 20711177 | Hamza TH | 08-15-2010 | Common genetic variation in the HLA region is associated with late-onset sporadic Parkinson's disease. | Nat Genet |
| Parkinson disease | 21084426 | Saad M | 11-17-2010 | Genome-wide association study confirms BST1 and suggests a locus on 12q24 as the risk loci for Parkin | Hum Mol Genet |
| Parkinson disease | 21248740 | Simón-Sánchez J | 01-19-2011 | Genome-wide association study confirms extant PD risk loci among the Dutch. | Eur J Hum Genet |
| Parkinson disease | 21292315 | Nalls MA | 02-01-2011 | Imputation of sequence variants for identification of genetic risks for Parkinson's disease: a meta-analy | Lancet |
| Parkinson disease | 21738487 | Do CB | 06-23-2011 | Web-based genome-wide association study identifies two novel loci and a substantial genetic compone | PLoS Genet |
| Parkinson disease | 21812969 | Liu X | 08-03-2011 | Genome-wide association study identifies candidate genes for Parkinson's disease in an Ashkenazi Jewi | BMC Med Genet |
| Parkinson disease | 22451204 | Pankratz N | 03-01-2012 | Meta-analysis of Parkinson's disease: identification of a novel locus, RIT2. | Ann Neurol |
| Parkinson disease | 22438815 | Lill CM | 03-15-2012 | Comprehensive research synopsis and systematic meta-analyses in Parkinson's disease genetics: The PI | PLoS Genet |
| Parkinson disease | 22911860 | Hernandez DG | 07-24-2012 | Genome wide assessment of young onset Parkinson's disease from Finland. | PLoS One |
| Parkinson disease | 24511991 | Hill-Burns EM | 02-10-2014 | Identification of a novel Parkinson's disease locus via stratified genome-wide association study. | BMC Genomics |
| Parkinson disease | 24842889 | Vacic V | 05-19-2014 | Genome-wide mapping of IBD segments in an Ashkenazi PD cohort identifies associated haplotypes. | Hum Mol Genet |
| Parkinson disease | 23793441 | Davis MF | 06-21-2013 | Parkinson disease loci in the mid-western Amish. | Hum Genet |
| Parkinson disease | 16252231 | Maraganore DM | 09-09-2005 | High-resolution whole-genome association study of Parkinson disease. | Am J Hum Genet |
| Parkinson disease | 19915576 | Satake W | 11-15-2009 | Genome-wide association study identifies common variants at four loci as genetic risk factors for Parkir | Nat Genet |
| Parkinson disease | 21044948 | Spencer CC | 11-02-2010 | Dissection of the genetics of Parkinson's disease identifies an additional association 5' of SNCA and mu | Hum Mol Genet |
| Parkinson disease | 22658654 | Chung SJ | 05-30-2012 | Genomic determinants of motor and cognitive outcomes in Parkinson's disease. | Parkinsonism Relat Disc |
| Parkinson disease | 25064009 | Nalls MA | 07-27-2014 | Large-scale meta-analysis of genome-wide association data identifies six new risk loci for Parkinson's c | Nat Genet |
| Parkinson disease | 25663231 | Beecham GW | 02-06-2015 | PARK10 is a major locus for sporadic neuropathologically confirmed Parkinson disease. | Neurology |
| Parkinson disease | 26227905 | Hu Y | 07-31-2015 | A Pooling Genome-Wide Association Study Combining a Pathway Analysis for Typical Sporadic Parkinso | Mol Neurobiol |
| Parkinson disease | 27545685 | Biernacka JM | 08-03-2016 | Genome-wide gene-environment interaction analysis of pesticide exposure and risk of Parkinson's disea | Parkinsonism Relat Disc |
| Parkinson disease | 28011712 | Foo JN | 12-22-2016 | Genome-wide association study of Parkinson's disease in East Asians. | Hum Mol Genet |
| Parkinson disease | 28892059 | Chang D | 09-11-2017 | A meta-analysis of genome-wide association studies identifies 17 new Parkinson's disease risk loci. | Nat Genet |
| Parkinson disease | 27182965 | Pickrell JK | 05-16-2016 | Detection and interpretation of shared genetic influences on 42 human traits. | Nat Genet |

| | | | | | |
|---|---|---|---|---|---|
| Parkinson disease | 30920090 | Trochet H | 03-28-2019 | Bayesian meta-analysis across genome-wide association studies of diverse phenotypes. | Genet Epidemiol |
| Parkinson disease | 26268663 | Nalls MA | 08-13-2015 | Baseline genetic associations in the Parkinson's Progression Markers Initiative (PPMI). | Mov Disord |
| Parkinson disease | 29724592 | Pottier C | 04-30-2018 | Potential genetic modifiers of disease risk and age at onset in patients with frontotemporal lobar degen | Lancet Neurol |
| Parkinson disease | 30338293 | Wallen ZD | 10-05-2018 | Plasticity-related gene 3 (LPPR1) and age at diagnosis of Parkinson disease. | Neurol Genet |
| Parkinson disease | 31701892 | Nalls MA | 12-01-2019 | Identification of novel risk loci, causal insights, and heritable risk for Parkinson's disease: a meta-analys | Lancet Neurol |
| Parkinson disease | 32310270 | Foo JN | 04-20-2020 | Identification of Risk Loci for Parkinson Disease in Asians and Comparison of Risk Between Asians and E | JAMA Neurol |
| Parkinson disease | 33111402 | Tan MMX | 10-28-2020 | Genome-Wide Association Studies of Cognitive and Motor Progression in Parkinson's Disease. | Mov Disord |
| Parkinson disease | 31660654 | Bandres-Ciga S | 10-29-2019 | The Genetic Architecture of Parkinson Disease in Spain: Characterizing Population-Specific Risk, Differen | Mov Disord |
| Parkinson disease | 32589924 | Bi W | 06-23-2020 | A Fast and Accurate Method for Genome-Wide Time-to-Event Data Analysis and Its Application to UK B | Am J Hum Genet |
| Parkinson disease | 31755958 | Blauwendraat C | 11-22-2019 | Genetic modifiers of risk and age at onset in GBA associated Parkinson's disease and Lewy body demer | Brain |
| Parkinson disease | 32201043 | Smeland OB | 02-08-2020 | Genome-wide Association Analysis of Parkinson's Disease and Schizophrenia Reveals Shared Genetic Ar | Biol Psychiatry |
| Parkinson disease | 33583074 | Le Guen Y | 03-06-2021 | Common X-Chromosome Variants Are Associated with Parkinson Disease Risk. | Ann Neurol |
| Parkinson disease | 34662886 | Backman JD | 10-18-2021 | Exome sequencing and analysis of 454,787 UK Biobank participants. | Nature |
| Parkinson disease | 34227697 | Loesch DP | 07-06-2021 | Characterizing the Genetic Architecture of Parkinson's Disease in Latinos. | Ann Neurol |
| Parkinson disease | 33987465 | Alfradique-Dunham | 01-28-2021 | Genome-Wide Association Study Meta-Analysis for Parkinson Disease Motor Subtypes. | Neurol Genet |
| Parkinson disease | 34064523 | Rodrigo LM | 05-04-2021 | Imputation and Reanalysis of ExomeChip Data Identifies Novel, Conditional and Joint Genetic Effects on | Genes (Basel) |
| Parkinson disease | 34776419 | Brolin K | 11-07-2021 | Insights on Genetic and Environmental Factors in Parkinson's Disease from a Regional Swedish Case-Co | J Parkinsons Dis |
| Parkinson disease | 34737426 | Jiang L | 11-04-2021 | A generalized linear mixed model association tool for biobank-scale data. | Nat Genet |
| Parkinson disease | 34594039 | Sakaue S | 09-30-2021 | A cross-population atlas of genetic associations for 220 human phenotypes. | Nat Genet |
| Alzheimer disease | 23565137 | Miyashita A | 04-02-2013 | SORL1 is genetically associated with late-onset Alzheimer's disease in Japanese, Koreans and Caucasiar | PLoS One |
| Alzheimer disease | 23571587 | Reitz C | 04-10-2013 | Variants in the ATP-binding cassette transporter (ABCA7), apolipoprotein E ϵ4,and the risk of late-onset | JAMA |
| Alzheimer disease | 23419831 | Ramanan VK | 02-19-2013 | APOE and BCHE as modulators of cerebral amyloid deposition: a florbetapir PET genome-wide associati | Mol Psychiatry |
| Alzheimer disease | 23562540 | Cruchaga C | 04-04-2013 | GWAS of cerebrospinal fluid tau levels identifies risk variants for Alzheimer's disease. | Neuron |
| Alzheimer disease | 23535033 | Sherva R | 03-24-2013 | Genome-wide association study of the rate of cognitive decline in Alzheimer's disease. | Alzheimers Dement |
| Alzheimer disease | 22832961 | Kamboh MI | 05-15-2012 | Genome-wide association study of Alzheimer's disease. | Transl Psychiatry |
| Alzheimer disease | 22881374 | Cummings AC | 09-01-2012 | Genome-wide association and linkage study in the Amish detects a novel candidate late-onset Alzheim | Ann Hum Genet |
| Alzheimer disease | 24755620 | Pérez-Palma E | 04-22-2014 | Overrepresentation of glutamate signaling in Alzheimer's disease: network-based pathway enrichment | PLoS One |
| Alzheimer disease | 24770881 | Nelson PT | 04-27-2014 | ABCC9 gene polymorphism is associated with hippocampal sclerosis of aging pathology. | Acta Neuropathol |
| Alzheimer disease | 23150908 | Jonsson T | 11-14-2012 | Variant of TREM2 associated with the risk of Alzheimer's disease. | N Engl J Med |
| Alzheimer disease | 23836404 | Shulman JM | 09-01-2013 | Genetic susceptibility for Alzheimer disease neuritic plaque pathology. | JAMA Neurol |
| Alzheimer disease | 18976728 | Bertram L | 10-29-2008 | Genome-wide association analysis reveals putative Alzheimer's disease susceptibility loci in addition to | Am J Hum Genet |
| Alzheimer disease | 19118814 | Beecham GW | 01-03-2009 | Genome-wide association study implicates a chromosome 12 risk locus for late-onset Alzheimer diseas | Am J Hum Genet |
| Alzheimer disease | 19734903 | Lambert JC | 09-06-2009 | Genome-wide association study identifies variants at CLU and CR1 associated with Alzheimer's disease | Nat Genet |
| Alzheimer disease | 20460622 | Seshadri S | 05-12-2010 | Genome-wide analysis of genetic loci associated with Alzheimer disease. | JAMA |
| Alzheimer disease | 20452100 | Kramer PL | 05-06-2010 | Alzheimer disease pathology in cognitively healthy elderly: a genome-wide study. | Neurobiol Aging |
| Alzheimer disease | 21059989 | Lee JH | 11-08-2010 | Identification of novel loci for Alzheimer disease and replication of CLU, PICALM, and BIN1 in Caribbean | Arch Neurol |
| Alzheimer disease | 17553421 | Reiman EM | 06-07-2007 | GAB2 alleles modify Alzheimer's risk in APOE epsilon4 carriers. | Neuron |
| Alzheimer disease | 17474819 | Coon KD | 04-01-2007 | A high-density whole-genome association study reveals that APOE is the major susceptibility gene for s | J Clin Psychiatry |
| Alzheimer disease | 25043464 | Jun G | 07-08-2014 | PLXNA4 is associated with Alzheimer disease and modulates tau phosphorylation. | Ann Neurol |
| Alzheimer disease | 22430674 | Lambert JC | 03-20-2012 | Genome-wide haplotype association study identifies the FRMD4A gene as a risk locus for Alzheimer's d | Mol Psychiatry |
| Alzheimer disease | 22005930 | Hollingworth P | 10-18-2011 | Genome-wide association study of Alzheimer's disease with psychotic symptoms. | Mol Psychiatry |
| Alzheimer disease | 21098978 | Sherva R | 11-25-2010 | Identification of novel candidate genes for Alzheimer's disease by autozygosity mapping using genome | J Alzheimers Dis |
| Alzheimer disease | 24162737 | Lambert JC | 10-27-2013 | Meta-analysis of 74,046 individuals identifies 11 new susceptibility loci for Alzheimer's disease. | Nat Genet |
| Alzheimer disease | 21460841 | Naj AC | 04-03-2011 | Common variants at MS4A4/MS4A6E, CD2AP, CD33 and EPHA1 are associated with late-onset Alzheime | Nat Genet |
| Alzheimer disease | 19125160 | Feulner TM | 01-07-2009 | Examination of the current top candidate genes for AD in a genome-wide association study. | Mol Psychiatry |

| | | | | | |
|---|---|---|---|---|---|
| Alzheimer disease | 22245343 | Meda SA | 01-08-2012 | A large scale multivariate parallel ICA method reveals novel imaging-genetic relationships for Alzheime | Neuroimage |
| Alzheimer disease | 22785395 | Gaj P | 07-11-2012 | Identification of a late onset Alzheimer's disease candidate risk variant at 9q21.33 in Polish patients. | J Alzheimers Dis |
| Alzheimer disease | 24958192 | Gusareva ES | 05-28-2014 | Genome-wide association interaction analysis for Alzheimer's disease. | Neurobiol Aging |
| Alzheimer disease | 18449908 | Poduslo SE | 04-30-2008 | Genome screen of late-onset Alzheimer's extended pedigrees identifies TRPC4AP by haplotype analysis | Am J Med Genet B Neu |
| Alzheimer disease | 21379329 | Wijsman EM | 02-17-2011 | Genome-wide association of familial late-onset Alzheimer's disease replicates BIN1 and CLU and nomi | PLoS Genet |
| Alzheimer disease | 25340798 | Kauwe JS | 10-23-2014 | Genome-wide association study of CSF levels of 59 alzheimer's disease candidate proteins: significant a | PLoS Genet |
| Alzheimer disease | 25649651 | Wang X | 02-03-2015 | Genetic Determinants of Survival in Patientswith Alzheimer’s Disease. | J Alzheimers Dis |
| Alzheimer disease | 26339675 | Tosto G | 06-18-2015 | F-box/LRR-repeat protein 7 is genetically associated with Alzheimer's disease. | Ann Clin Transl Neurol |
| Alzheimer disease | 25778476 | Jun G | 03-17-2015 | A novel Alzheimer disease locus located near the gene encoding tau protein. | Mol Psychiatry |
| Alzheimer disease | 21116278 | Furney SJ | 11-30-2010 | Genome-wide association with MRI atrophy measures as a quantitative trait locus for Alzheimer's disea | Mol Psychiatry |
| Alzheimer disease | 26049409 | Hirano A | 06-05-2015 | A genome-wide association study of late-onset Alzheimer's disease in a Japanese population. | Psychiatr Genet |
| Alzheimer disease | 26913989 | Traylor M | 02-23-2016 | Shared genetic contribution to Ischaemic Stroke and Alzheimer's Disease. | Ann Neurol |
| Alzheimer disease | 28183528 | Jun GR | 02-06-2017 | Transethnic genome-wide scan identifies novel Alzheimer's disease loci. | Alzheimers Dement |
| Alzheimer disease | 28560309 | Lee E | 04-23-2017 | Single-nucleotide polymorphisms are associated with cognitive decline at Alzheimer's disease conversio | Alzheimers Dement (Ar |
| Alzheimer disease | 28870582 | Wang XF | 08-01-2017 | Linking Alzheimer's disease and type 2 diabetes: Novel shared susceptibility genes detected by cFDR ap | J Neurol Sci |
| Alzheimer disease | 26993346 | Schott JM | 03-15-2016 | Genetic risk factors for the posterior cortical atrophy variant of Alzheimer's disease. | Alzheimers Dement |
| Alzheimer disease | 30636644 | Nazarian A | 01-12-2019 | Genome-wide analysis of genetic predisposition to Alzheimer's disease and related sex disparities. | Alzheimers Res Ther |
| Alzheimer disease | 30413934 | Broce IJ | 11-09-2018 | Dissecting the genetic relationship between cardiovascular risk factors and Alzheimer's disease. | Acta Neuropathol |
| Alzheimer disease | 31473137 | Moreno-Grau S | 08-13-2019 | Genome-wide association analysis of dementia and its clinical endophenotypes reveal novel loci associa | Alzheimers Dement |
| Alzheimer disease | 29615537 | Kulminski AM | 03-01-2018 | Strong impact of natural-selection-free heterogeneity in genetics of age-related phenotypes. | Aging (Albany NY) |
| Alzheimer disease | 29777097 | Marioni RE | 05-18-2018 | GWAS on family history of Alzheimer's disease. | Transl Psychiatry |
| Alzheimer disease | 30617256 | Jansen IE | 01-07-2019 | Genome-wide meta-analysis identifies new loci and functional pathways influencing Alzheimer's diseas | Nat Genet |
| Alzheimer disease | 31055733 | Nazarian A | 05-05-2019 | Genetic heterogeneity of Alzheimer's disease in subjects with and without hypertension. | Geroscience |
| Alzheimer disease | 30514930 | Mukherjee S | 12-04-2018 | Genetic data and cognitively defined late-onset Alzheimer's disease subgroups. | Mol Psychiatry |
| Alzheimer disease | 30805717 | Zhu Z | 02-25-2019 | Shared genetic architecture between metabolic traits and Alzheimer's disease: a large-scale genome-w | Hum Genet |
| Alzheimer disease | 30201328 | Gusareva ES | 08-09-2018 | Male-specific epistasis between WWC1 and TLN2 genes is associated with Alzheimer's disease. | Neurobiol Aging |
| Alzheimer disease | 29107063 | Yashin AI | 10-26-2017 | Hidden heterogeneity in Alzheimer's disease: Insights from genetic association studies and other analys | Exp Gerontol |
| Alzheimer disease | 30979435 | Lo MT | 03-12-2019 | Identification of genetic heterogeneity of Alzheimer's disease across age. | Neurobiol Aging |
| Alzheimer disease | 32589924 | Bi W | 06-23-2020 | A Fast and Accurate Method for Genome-Wide Time-to-Event Data Analysis and Its Application to UK B | Am J Hum Genet |
| Alzheimer disease | 32641180 | Steffens DC | 07-09-2020 | Genome-wide screen to identify genetic loci associated with cognitive decline in late-life depression. | Int Psychogeriatr |
| Alzheimer disease | 33541420 | Bone WP | 02-04-2021 | Multi-trait association studies discover pleiotropic loci between Alzheimer's disease and cardiometabol | Alzheimers Res Ther |
| Alzheimer disease | 32588970 | Lutz MW | 06-26-2020 | Shared genetic etiology underlying late-onset Alzheimer's disease and posttraumatic stress syndrome. | Alzheimers Dement |
| Alzheimer disease | 33589840 | Schwartzentruber J | 02-15-2021 | Genome-wide meta-analysis, fine-mapping and integrative prioritization implicate new Alzheimer's dis | Nat Genet |
| Alzheimer disease | 33223526 | Hong S | 11-22-2020 | Genome-wide association study of Alzheimer's disease CSF biomarkers in the EMIF-AD Multimodal Bio | Transl Psychiatry |
| Alzheimer disease | 33654092 | Shigemizu D | 03-03-2021 | Ethnic and trans-ethnic genome-wide association studies identify new loci influencing Japanese Alzheir | Transl Psychiatry |
| Alzheimer disease | 33074286 | Kunkle BW | 10-19-2020 | Novel Alzheimer Disease Risk Loci and Pathways in African American Individuals Using the African Genc | JAMA Neurol |
| Alzheimer disease | 34122051 | Wang H | 05-28-2021 | Similar Genetic Architecture of Alzheimer's Disease and Differential <i>APOE</i> Effect Between Sexes | Front Aging Neurosci |
| Alzheimer disease | 34011927 | Park JH | 05-19-2021 | Novel Alzheimer's disease risk variants identified based on whole-genome sequencing of APOE ε4 carri | Transl Psychiatry |
| Alzheimer disease | 34099642 | de Rojas I | 06-07-2021 | Common variants in Alzheimer's disease and risk stratification by polygenic risk scores. | Nat Commun |
| Alzheimer disease | 34336000 | Li L | 07-14-2021 | Use of Deep-Learning Genomics to Discriminate Healthy Individuals from Those with Alzheimer's Disea | Behav Neurol |
| Alzheimer disease | 34743297 | Kulminski AM | 11-06-2021 | Pleiotropic predisposition to Alzheimer's disease and educational attainment: insights from the summa | Geroscience |
| Alzheimer disease | 35379992 | Bellenguez C | 04-04-2022 | New insights into the genetic etiology of Alzheimer's disease and related dementias. | Nat Genet |
| Alzheimer disease | 34662886 | Backman JD | 10-18-2021 | Exome sequencing and analysis of 454,787 UK Biobank participants. | Nature |
| Alzheimer disease | 35770850 | Chung J | 06-30-2022 | Genome-wide association and multi-omics studies identify MGMT as a novel risk gene for Alzheimer's | Alzheimers Dement |
| Alzheimer disease | 35589863 | Gouveia C | 05-19-2022 | Genome-wide association of polygenic risk extremes for Alzheimer's disease in the UK Biobank. | Sci Rep |

| | | | | | |
|---|---|---|---|---|---|
| Alzheimer disease | 35879306 | Hou J | 07-25-2022 | Polygenic resilience scores capture protective genetic effects for Alzheimer's disease. | Transl Psychiatry |
| Alzheimer disease | 35851147 | Adewuyi EO | 07-18-2022 | A large-scale genome-wide cross-trait analysis reveals shared genetic architecture between Alzheimer' | Commun Biol |
| Alzheimer disease | 35977442 | Bracher-Smith M | 07-29-2022 | Whole genome analysis in APOE4 homozygotes identifies the DAB1-RELN pathway in Alzheimer's disea | Neurobiol Aging |

**Table S2: Overview of synaptic suggestive GWAS candidate genes associated with Alzheimer disease, Parkinson disease, schizophrenia and autism spectrum disorder.**

| Disease | Gene symbol | Gene description |
|---|---|---|
| Autism spectrum disorder | AGER | Advanced glycosylation end-product specific receptor |
| Autism spectrum disorder | ANK3 | Ankyrin 3 |
| Autism spectrum disorder | ANKS1B | Ankyrin repeat and sterile alpha motif domain containing 1B |
| Autism spectrum disorder | ATP6V1G2 | ATPase H+ transporting V1 subunit G2 |
| Autism spectrum disorder | C4A | Complement C4A |
| Autism spectrum disorder | C4B | Complement C4B |
| Autism spectrum disorder | C4B_2 | Complement component 4B, copy 2 |
| Autism spectrum disorder | CACNA1C | Calcium voltage-gated channel subunit alpha1 C |
| Autism spectrum disorder | CACNB2 | Calcium voltage-gated channel auxiliary subunit beta 2 |
| Autism spectrum disorder | CHRM4 | Cholinergic receptor muscarinic 4 |
| Autism spectrum disorder | CHRNA3 | Cholinergic receptor nicotinic alpha 3 subunit |
| Autism spectrum disorder | CHRNA5 | Cholinergic receptor nicotinic alpha 5 subunit |
| Autism spectrum disorder | CHRNB4 | Cholinergic receptor nicotinic beta 4 subunit |
| Autism spectrum disorder | CLU | Clusterin |
| Autism spectrum disorder | CSNK2B | Casein kinase 2 beta |
| Autism spectrum disorder | CTNND1 | Catenin delta 1 |
| Autism spectrum disorder | DGKI | Diacylglycerol kinase iota |
| Autism spectrum disorder | DGKZ | Diacylglycerol kinase zeta |
| Autism spectrum disorder | DRD2 | Dopamine receptor D2 |
| Autism spectrum disorder | FLOT1 | Flotillin 1 |
| Autism spectrum disorder | GABBR1 | Gamma-aminobutyric acid type B receptor subunit 1 |
| Autism spectrum disorder | GRIN2A | Glutamate ionotropic receptor NMDA type subunit 2A |
| Autism spectrum disorder | GRM3 | Glutamate metabotropic receptor 3 |
| Autism spectrum disorder | HCN1 | Hyperpolarization activated cyclic nucleotide gated potassiu |
| Autism spectrum disorder | IGSF9B | Immunoglobulin superfamily member 9B |
| Autism spectrum disorder | LRP4 | LDL receptor related protein 4 |
| Autism spectrum disorder | LY6G6D | Lymphocyte antigen 6 family member G6D |
| Autism spectrum disorder | LY6G6E | Lymphocyte antigen 6 family member G6E (pseudogene) |
| Autism spectrum disorder | MEF2C | Myocyte enhancer factor 2C |
| Autism spectrum disorder | MOB4 | MOB family member 4, phocein |
| Autism spectrum disorder | NGEF | Neuronal guanine nucleotide exchange factor |
| Autism spectrum disorder | NRGN | Neurogranin |
| Autism spectrum disorder | PRRT1 | Proline rich transmembrane protein 1 |
| Autism spectrum disorder | PTPRF | Protein tyrosine phosphatase receptor type F |
| Autism spectrum disorder | RIMS1 | Regulating synaptic membrane exocytosis 1 |
| Autism spectrum disorder | SCGN | Secretagogin, EF-hand calcium binding protein |
| Autism spectrum disorder | SNAP91 | Synaptosome associated protein 91 |
| Autism spectrum disorder | SORCS3 | Sortilin related VPS10 domain containing receptor 3 |
| Autism spectrum disorder | SYNGAP1 | Synaptic Ras GTPase activating protein 1 |
| Autism spectrum disorder | ZNF804A | Zinc finger protein 804A |
| Schizophrenia | ADAM10 | ADAM metallopeptidase domain 10 |
| Schizophrenia | AGER | Advanced glycosylation end-product specific receptor |
| Schizophrenia | ANK3 | Ankyrin 3 |
| Schizophrenia | ANKS1B | Ankyrin repeat and sterile alpha motif domain containing 1B |
| Schizophrenia | ATP2A2 | ATPase sarcoplasmic/endoplasmic reticulum Ca2+ transport |

| | | |
|---|---|---|
| Schizophrenia | ATP6V1G2 | ATPase H+ transporting V1 subunit G2 |
| Schizophrenia | C4A | Complement C4A |
| Schizophrenia | C4B | Complement C4B |
| Schizophrenia | C4B_2 | Complement component 4B, copy 2 |
| Schizophrenia | CACNA1C | Calcium voltage-gated channel subunit alpha1 C |
| Schizophrenia | CACNB2 | Calcium voltage-gated channel auxiliary subunit beta 2 |
| Schizophrenia | CDH13 | Cadherin 13 |
| Schizophrenia | CHRM4 | Cholinergic receptor muscarinic 4 |
| Schizophrenia | CHRNA2 | Cholinergic receptor nicotinic alpha 2 subunit |
| Schizophrenia | CHRNA3 | Cholinergic receptor nicotinic alpha 3 subunit |
| Schizophrenia | CHRNA5 | Cholinergic receptor nicotinic alpha 5 subunit |
| Schizophrenia | CHRNB4 | Cholinergic receptor nicotinic beta 4 subunit |
| Schizophrenia | CLCN3 | Chloride voltage-gated channel 3 |
| Schizophrenia | CLU | Clusterin |
| Schizophrenia | CNTN2 | Contactin 2 |
| Schizophrenia | CPEB1 | Cytoplasmic polyadenylation element binding protein 1 |
| Schizophrenia | CSNK2B | Casein kinase 2 beta |
| Schizophrenia | CTNND1 | Catenin delta 1 |
| Schizophrenia | DCC | DCC netrin 1 receptor |
| Schizophrenia | DGKI | Diacylglycerol kinase iota |
| Schizophrenia | DGKZ | Diacylglycerol kinase zeta |
| Schizophrenia | DLG2 | Discs large MAGUK scaffold protein 2 |
| Schizophrenia | DOC2A | Double C2 domain alpha |
| Schizophrenia | DRD2 | Dopamine receptor D2 |
| Schizophrenia | DSCAM | DS cell adhesion molecule |
| Schizophrenia | EEF1D | Eukaryotic translation elongation factor 1 delta |
| Schizophrenia | ELAVL2 | ELAV like RNA binding protein 2 |
| Schizophrenia | EMB | Embigin |
| Schizophrenia | FLOT1 | Flotillin 1 |
| Schizophrenia | FXR1 | FMR1 autosomal homolog 1 |
| Schizophrenia | FYN | FYN proto-oncogene, Src family tyrosine kinase |
| Schizophrenia | GABBR1 | Gamma-aminobutyric acid type B receptor subunit 1 |
| Schizophrenia | GABBR2 | Gamma-aminobutyric acid type B receptor subunit 2 |
| Schizophrenia | GPM6A | Glycoprotein M6A |
| Schizophrenia | GRIA1 | Glutamate ionotropic receptor AMPA type subunit 1 |
| Schizophrenia | GRIK3 | Glutamate ionotropic receptor kainate type subunit 3 |
| Schizophrenia | GRIN2A | Glutamate ionotropic receptor NMDA type subunit 2A |
| Schizophrenia | GRM3 | Glutamate metabotropic receptor 3 |
| Schizophrenia | HAPLN4 | Hyaluronan and proteoglycan link protein 4 |
| Schizophrenia | HCN1 | Hyperpolarization activated cyclic nucleotide gated potassiu |
| Schizophrenia | IGSF9B | Immunoglobulin superfamily member 9B |
| Schizophrenia | INA | Internexin neuronal intermediate filament protein alpha |
| Schizophrenia | KCNB1 | Potassium voltage-gated channel subfamily B member 1 |
| Schizophrenia | LPAR2 | Lysophosphatidic acid receptor 2 |
| Schizophrenia | LRP4 | LDL receptor related protein 4 |
| Schizophrenia | LRRTM4 | Leucine rich repeat transmembrane neuronal 4 |
| Schizophrenia | LY6G6D | Lymphocyte antigen 6 family member G6D |
| Schizophrenia | LY6G6E | Lymphocyte antigen 6 family member G6E (pseudogene) |

| | | |
|---|---|---|
| Schizophrenia | MAGI2 | Membrane associated guanylate kinase, WW and PDZ doma |
| Schizophrenia | MEF2C | Myocyte enhancer factor 2C |
| Schizophrenia | MOB4 | MOB family member 4, phocein |
| Schizophrenia | NCAN | Neurocan |
| Schizophrenia | NGEF | Neuronal guanine nucleotide exchange factor |
| Schizophrenia | NLGN4X | Neuroligin 4 X-linked |
| Schizophrenia | NOS1 | Nitric oxide synthase 1 |
| Schizophrenia | NRGN | Neurogranin |
| Schizophrenia | OLFM3 | Olfactomedin 3 |
| Schizophrenia | PCDHA4 | Protocadherin alpha 4 |
| Schizophrenia | PCLO | Piccolo presynaptic cytomatrix protein |
| Schizophrenia | PDE4B | Phosphodiesterase 4B |
| Schizophrenia | PRR12 | Proline rich 12 |
| Schizophrenia | PRRT1 | Proline rich transmembrane protein 1 |
| Schizophrenia | PSD3 | Pleckstrin and Sec7 domain containing 3 |
| Schizophrenia | PTK2B | Protein tyrosine kinase 2 beta |
| Schizophrenia | PTN | Pleiotrophin |
| Schizophrenia | PTPRF | Protein tyrosine phosphatase receptor type F |
| Schizophrenia | RIMS1 | Regulating synaptic membrane exocytosis 1 |
| Schizophrenia | SCGN | Secretagogin, EF-hand calcium binding protein |
| Schizophrenia | SHISA8 | Shisa family member 8 |
| Schizophrenia | SLC12A4 | Solute carrier family 12 member 4 |
| Schizophrenia | SLC32A1 | Solute carrier family 32 member 1 |
| Schizophrenia | SLC4A10 | Solute carrier family 4 member 10 |
| Schizophrenia | SNAP91 | Synaptosome associated protein 91 |
| Schizophrenia | SORCS3 | Sortilin related VPS10 domain containing receptor 3 |
| Schizophrenia | STAC3 | SH3 and cysteine rich domain 3 |
| Schizophrenia | SYNGAP1 | Synaptic Ras GTPase activating protein 1 |
| Schizophrenia | TAOK2 | TAO kinase 2 |
| Schizophrenia | TENM2 | Teneurin transmembrane protein 2 |
| Schizophrenia | VPS45 | Vacuolar protein sorting 45 homolog |
| Schizophrenia | YWHAE | Tyrosine 3-monooxygenase/tryptophan 5-monooxygenase a( |
| Schizophrenia | ZDHHC2 | Zinc finger DHHC-type palmitoyltransferase 2 |
| Schizophrenia | ZDHHC5 | zinc finger DHHC-type palmitoyltransferase 5 |
| Schizophrenia | ZNF804A | Zinc finger protein 804A |
| Parkinson disease | APOE | Apolipoprotein E |
| Parkinson disease | CAMK2D | Calcium/calmodulin dependent protein kinase II delta |
| Parkinson disease | DGKQ | Diacylglycerol kinase theta |
| Parkinson disease | DLG2 | Discs large MAGUK scaffold protein 2 |
| Parkinson disease | FYN | FYN proto-oncogene, Src family tyrosine kinase |
| Parkinson disease | GAK | Cyclin G associated kinase |
| Parkinson disease | HIP1R | Huntingtin interacting protein 1 related |
| Parkinson disease | IGSF9B | Immunoglobulin superfamily member 9B |
| Parkinson disease | ITGA8 | Integrin subunit alpha 8 |
| Parkinson disease | LRRK2 | Leucine rich repeat kinase 2 |
| Parkinson disease | MAPT | Microtubule associated protein tau |

| | | |
|---|---|---|
| Parkinson disease | MCTP2 | Multiple C2 and transmembrane domain containing 2 |
| Parkinson disease | NSF | N-ethylmaleimide sensitive factor, vesicle fusing ATPase |
| Parkinson disease | RIT2 | Ras like without CAAX 2 |
| Parkinson disease | SH3GL2 | SH3 domain containing GRB2 like 2, endophilin A1 |
| Parkinson disease | SNCA | Synuclein alpha |
| Parkinson disease | STX1B | Syntaxin 1B |
| Parkinson disease | SYT11 | Synaptotagmin 11 |
| Parkinson disease | SYT4 | Synaptotagmin 4 |
| Parkinson disease | TMEM163 | Transmembrane protein 163 |
| Alzheimer disease | ADAM10 | ADAM metallopeptidase domain 10 |
| Alzheimer disease | AP2A2 | Adaptor related protein complex 2 subunit alpha 2 |
| Alzheimer disease | APOE | Apolipoprotein E |
| Alzheimer disease | BIN1 | Bridging integrator 1 |
| Alzheimer disease | CLU | Clusterin |
| Alzheimer disease | CNTNAP2 | Contactin associated protein 2 |
| Alzheimer disease | GPC6 | Glypican 6 |
| Alzheimer disease | MAPT | Microtubule associated protein tau |
| Alzheimer disease | MEF2C | Myocyte enhancer factor 2C |
| Alzheimer disease | NCK2 | NCK adaptor protein 2 |
| Alzheimer disease | PICALM | Phosphatidylinositol binding clathrin assembly protein |
| Alzheimer disease | PTK2B | Protein tyrosine kinase 2 beta |
| Alzheimer disease | TLN2 | Talin 2 |
